# Filamented Light (FLight) Bioprinting of Mini-Muscles with Self-Renewal Potential

**DOI:** 10.1101/2025.02.03.636126

**Authors:** Hao Liu, Michael Winkelbauer, Jakub Janiak, Ali Kerem Kalkan, Inseon Kim, Parth Chansoria, Ori Bar-Nur, Marcy Zenobi-Wong

## Abstract

The plasticity and regenerative capacity of skeletal muscle arise from quiescent stem cells activated upon overload, injury, or inflammation. Developing *in vitro* muscle models to study these properties could advance muscle disease modeling and pre-clinical evaluation. Here, we leverage Filamented Light (FLight) bioprinting as a high-throughput approach for producing mini-muscle tissues. Using Pax7-nGFP myoblasts, we bioprinted mini-muscles from pristine collagen-fibrinogen. The FLight hydrogel consisted of aligned microstructures which guided the formation of aligned myotubes. Mini-muscles demonstrated *in vivo*-like tissue organization, including highly aligned myotubes and a Pax7^+^ cell pool embedded in newly deposited laminin. Both spontaneous and electrically stimulated contractions were observed. Collagen-fibrinogen matrix was promising for maintenance of the Pax7^+^ cell pool. Damage from cardiotoxin-induced injury of the mini-muscles led to a massive proliferation of Pax7^+^ cells and restoration of the contractile properties. Notably, small molecules such as Repsox could enhance regeneration. FLight printed mini-muscles have potential for applications in muscle biology, exercise/atrophy, disease models, and drug screening.

## Introduction

Skeletal muscles have remarkable plasticity, adapting their metabolism and tissue structure in response to environmental signals, such as mechanical loading.[1] This plasticity is most apparent in muscle’s remarkable regeneration capacity upon injury.[2] Muscle regeneration is primarily driven by a muscle-resident population of satellite cells with myogenic stem cells function. These satellite cells reside within a stem cell niche and are maintained in a quiescent state through essential signaling pathways, such as the Notch and integrin pathways.[3–5] Satellite cells are commonly identified through expression of the transcription factor *paired homeobox protein 7* (Pax7), which regulates the myogenic pathway. Upon muscle injury, a subset of quiescent Pax7^+^ cells re-enter the cell cycle, become myogenic, and ultimately fuse with damaged muscle fibers to help restore muscle function.

Satellite cells cultured *in vitro* in 2D monolayers often lose Pax7 expression, primarily due to the absence of the 3D niche environment and associated signaling pathways.[5] *In vivo*, the fate of satellite cells is tightly regulated by their stem cell niche, which consists of two critical elements: the host muscle fiber and the basal lamina.[6] Mechanical and chemical signals from the host muscle fiber directly influence satellite cell behavior, such as activation and proliferation,[7] while the basal lamina provides the necessary matrix microenvironment. These components work together to regulate satellite cell quiescence, activation, and proliferation.

The basal lamina, a key extracellular matrix (ECM) component in muscle tissue, is primarily composed of laminin, collagen, and proteoglycans.[8,9] Satellite cells are anchored to the basal lamina through cell-matrix interactions (e.g., integrin family),[8] which are critical for maintaining their stem cell state. For instance, previous studies have shown that satellite cells not only express Pax7, but also cell surface markers such as integrin beta 1 (ITGB1).[5] In fact, the absence of ITGB1 leads to an inability of satellite cells to maintain their quiescence and depletion of the satellite cell pool.[10] This presents a significant challenge for long-term *in vitro* culture and muscle tissue engineering using myogenic stem cells.

There is an urgent need for engineered *in vitro* muscle models to study the role of myogenic stem cells in muscle physiology and pathology. Such engineered muscle models should mimic key attributes of skeletal muscle tissues: highly aligned muscle fibers, contractility, and self-renewal potential mediated by myogenic stem cells. Tissue engineering strategies should provide effective topological cues to support stem cell differentiation into aligned muscle fibers and should emulate the *in vivo* tissue environment to maintain the stem cell pool. Perhaps the most common approach for fabricating *in vitro* muscle tissues involves casting collagen, fibrinogen, Matrigel™, or a combination of these materials between flexible PDMS posts. During culture and differentiation, the cells remodel the matrix and fuse, resulting in aligned, multinucleated myotubes that exhibit several features of mature muscle, including sarcomere structures and myosin heavy chain staining, as well as the presence of extracellular laminin and collagen I. These tissue constructs have been successfully used for disease modeling and drug testing.[11,12]

However, there are notable limitations that hinder the versatility and repeatability of the casting approach. Aligned myotubes and multinucleation seen in the casting method are largely dependent on the cells’ ability to deform and remodel the matrix, which can be affected by the cell type or donor-to-donor variability. For example, matrix deformation, and consequently muscle alignment and maturation, may differ due to the lower reproducibility often associated with the casting approach. These factors introduce variability in tissue models and limit their use in preclinical drug testing. There have been alternative approaches, such as extrusion-based bioprinting, surface patterning, acoustic patterning, and electrospinning, which rely on topological cues to orient myotubes. However, these methods still require substantial exploration to develop muscle tissues comparable to the casting method.[13–16] Overall, only a limited number of studies have reported the *in vitro* generation of functional muscle from primary myogenic stem cells, especially with self-renewal potential.[17]

Filamented light (FLight) bioprinting allows for the biofabrication of many aligned cell-laden anisotropic hydrogels within seconds.[18] The inhomogeneous distribution of light intensity in the laser beam, caused by the optical modulation instability (OMI) effect, results in **1.** highly aligned microfilaments with diameters ranging from 2 to 20 µm, which serve as topological cues, and **2.** interconnected channel-like voids with diameters ranging from 2 to 20 µm, known as microchannels. In our previous studies, we have shown that ultra-high aspect-ratio gelatin microstructures support the proliferation and fusion of C2C12 cells into contractile myotubes.

Recent advances in the development of photocrosslinkable biomaterials have expanded the potential of light-based biofabrication, including the use of pristine proteins. For example, it has been demonstrated that ruthenium (Ru) and sodium persulfate (SPS) can be used as photoinitiators to crosslink collagen type I, fibrinogen, silk fibroin, and decellularized extracellular matrix (dECM) under visible light.[19–22] In these materials, the Ru-SPS initiator system facilitated photocrosslinking of native tyrosine residues in the materials, eliminating the need for chemical modification. This approach allows the unmodified biomaterials to retain their natural properties and bioactivity, such as enabling integrin-mediated cell adhesion.[23] However, the solubility of collagen type I is highly dependent on the pH of the solvent. Collagen solutions are often prepared in an acidic environment (pH 2-5), to prevent self-assembly of collagen fibrils (also known as fibrillogenesis),[24] which also protects against the physical crosslinking of collagen into a hydrogel at a macroscopic level.[25] However, the acidic environment significantly hinders the application of collagen in tissue engineering and biofabrication.

Here, we report on the use of FLight biofabrication of pristine collagen and fibrinogen to create filamented hydrogel matrices under neutral conditions (**Figure 1a**). Compared to conventional gel casting via thermal and enzymatic crosslinking of collagen and fibrinogen,[26–28] light-based printing allows for the creation of customized hydrogel structures containing internal microstructures. The pristine protein-based FLight matrices not only facilitate the fusion of mouse Pax7-nGFP primary myoblasts into aligned myotubes but also better maintain the Pax7^+^ cell pool compared to norbornene-thiol-functionalized gelatin matrices. After 3 weeks of differentiation, the engineered FLight muscle exhibited tissue organization similar to that observed *in vivo*, along with spontaneous contractions. Notably, the engineered FLight muscle demonstrated self-renewal capacity, which could be further enhanced by small molecules such as Forskolin, Chir 99021, and Repsox. This engineered muscle model holds great potential as an *in vitro* screening platform to accelerate the development of novel therapies for skeletal muscle diseases.

**Figure 1.**
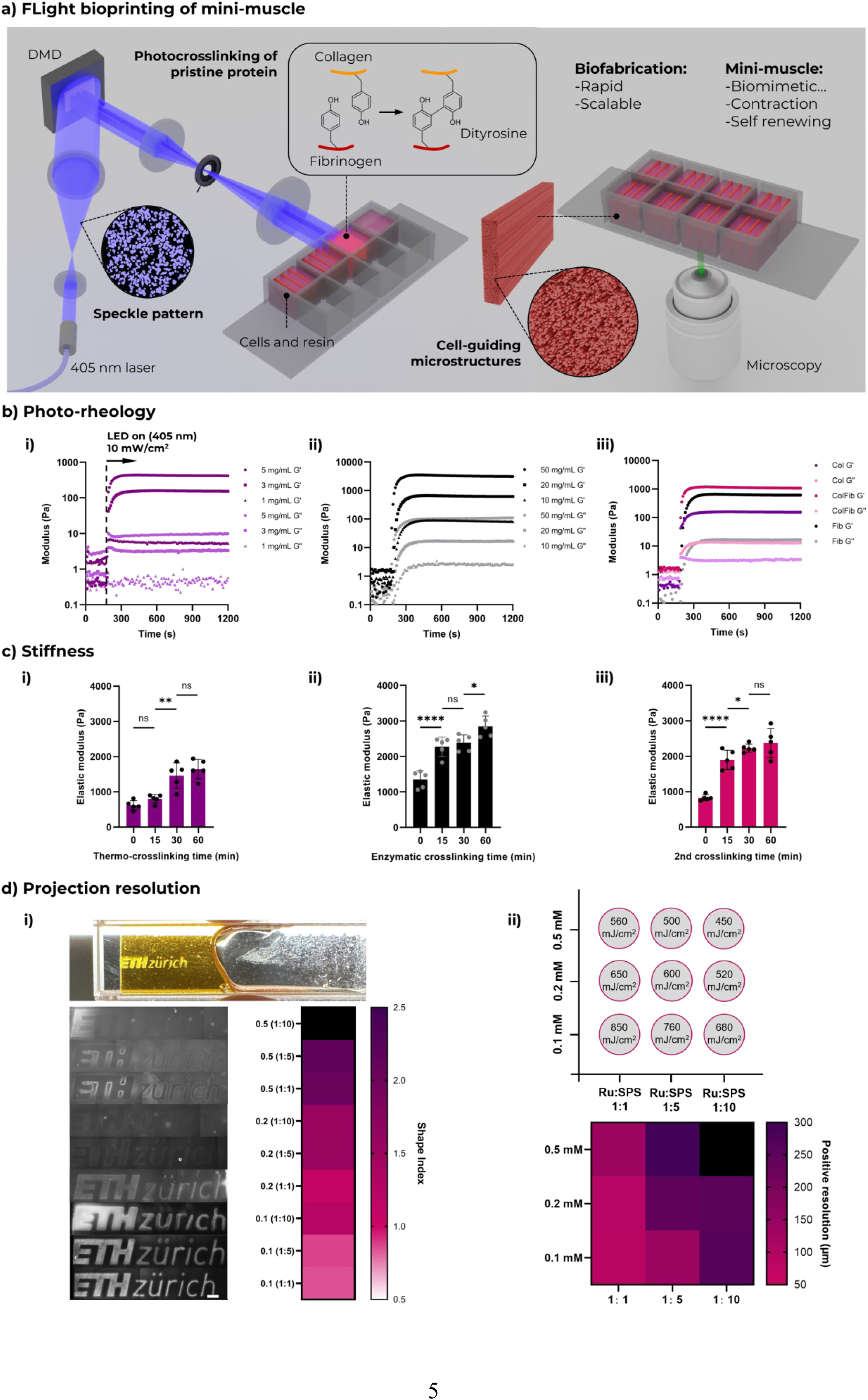
Photocrosslinking of pristine Collagen/ Fibrinogen and Its Application in FLight Printing. **a)** Schematic illustration of FLight printing of engineered mini-muscles using pristine protein-based photoresins. **b**) Rheological analysis of photocrosslinking for i) pure collagen (Col), ii) pure fibrinogen (Fib) and iii) their mixture (3 mg/mL Col + 25 mg/mL Fib; compared to 3 mg/mL Col and 20 mg/mL Fib) at different protein concentrations under 405 nm wavelengths of light, using ruthenium-SPS as the photoinitiator. **c**) Compressive elastic modulus of FLight hydrogel constructs before and after secondary crosslinking. i) Hydrogels printed with 5 mg/mL pure collagen photoresin, measured after 0, 15, 30, and 60 minutes of thermal crosslinking at 37 °C. ii) Elastic modulus of hydrogels printed with fibrinogen photoresin at a concentration of 50 mg/mL, following 15, 30, and 60 minutes of enzymatic crosslinking using thrombin at 37 °C. iii) Elastic modulus of hydrogel samples printed with ColFib photoresin (3 mg/mL Col + 25 mg/mL Fib), and after 15, 30, and 60 minutes of secondary crosslinking at 37 °C. **d**) Resolution assessments for FLight printing using ColFib photoresin with varying concentrations/ratios of Ru-SPS photoinitiator (e.g., 0.5 (1:10) refers to 0.5 mM Ru + 5 mM SPS). i) Evaluation of printing fidelity by comparing the designed feature size to the printed feature size (shape index). A shape index closest to 1.0 indicates optimal printing fidelity. Scale bar: 500 µm. ii) Matrix of light dose used in the resolution test and the corresponding minimum positive feature size (positive resolution). Data in c) are presented as mean ± SEM and individual points (*N* = 5); \**P* < 0.05, \*\**P* < 0.01, \*\*\**P* < 0.001, and \*\*\*\**P* < 0.0001.

## Results

### Photocrosslinking of pristine collagen, fibrinogen and their mixture

The self-assembly of collagen fibrils requires the formation of hydrogen-bonded water clusters,[29] and the glucose solution competes with water molecules for the hydrogen bonds critical for fibrillogenesis,[30] thereby temporarily preventing the thermal crosslinking of collagen. No temperature-dependent crosslinking was detected when collagen type I was dissolved in a glucose-phosphate-rich buffer system which temporarily inhibits the fibrillogenic potential of collagen (pH 7.4, **Figure S1**). However, thermal crosslinking was observed when collagen was dissolved in a phosphate-buffered saline (PBS) solution. As the hydrogen bonds in the collagen fibrillogenesis inhibition (CFI) buffer were not permanent, the collagen solution could crosslink once the CFI was replaced with PBS solution (**Figure S2**). We then confirmed with circular dichroic spectroscopy that the native collagen type I molecules could fold into a triple-helix, where a positive peak at ≈ 220 nm indicated triple helical structure in different solutions (acetic acid, PBS, and CFI) (**Figure S3**). This approach offers a broad biofabrication window for pristine collagen that is both temperature- and pH-insensitive, while preserving the unique triple-helix structure of collagen type I, thereby maximizing its bioactivity and functionality in various tissue engineering applications.

We then evaluated the photocrosslinking potential of pristine proteins, including collagen, fibrinogen, and their mixtures, using ruthenium-sodium persulfate (Ru-SPS) as a photoinitiator through photorheological testing (**Figure 1b**). With a final concentration of 0.2 mM Ru and 2 mM SPS in various protein-based photoresins, an increase in storage modulus was observed upon exposure to 405 nm wavelength LED (∼ 10 mW/cm^2^). The storage modulus plateaued within 120 seconds for all protein-based photoresins, indicating successful photocrosslinking. Notably, although the final storage modulus varied depending on the initial protein concentration, pure collagen-based photoresin exhibited lower mechanical properties compared to fibrinogen. Increasing the collagen type I concentration could enhance the stiffness of the final hydrogel constructs, but excessive rigidity can make such photoresins difficult to work with. Therefore, a mixture of collagen and fibrinogen was selected as the optimal formulation for subsequent studies.

Despite the ideal photocrosslinking kinetics of the protein-based photoresins, the stiffness of the resulting hydrogel constructs was insufficient for culture and applications. Therefore, to enhance the mechanical properties of the hydrogels, we tested a secondary crosslinking strategy based on the properties of individual proteins, including thermo-crosslinking of collagen and enzymatic crosslinking of fibrinogen using thrombin (**Figure 1c**). The elastic modulus of photocrosslinked collagen hydrogel constructs increased after replacing the fibrillogenesis inhibition buffer with PBS and incubating at 37°C (light dose: 600 mJ/cm²). The mean compressive stiffness of the pure collagen hydrogels reached 1.7 kPa after 60 minutes of incubation, compared to approximately 0.6 kPa immediately following photocrosslinking. This suggests that a secondary crosslinking process, analogous to thermal crosslinking in a neutral environment, had occurred.

Additionally, thrombin, an enzyme that facilitates the polymerization of fibrinogen into fibrin, was used as a secondary crosslinking strategy for fibrinogen-based hydrogels. Following photocrosslinking (light dose: 600 mJ/cm²), the hydrogel samples were incubated in PBS containing 5 U/mL of thrombin at 37°C. The stiffness of the pure fibrinogen hydrogels increased from an initial value of 1.3 kPa (without enzymatic crosslinking) to 2.8 kPa. For ColFib hydrogels (i.e., those having both collagen Type I and fibrinogen), incubation with thrombin at 37°C enabled the simultaneous secondary crosslinking of both photocrosslinked collagen and fibrinogen. Compared to the initial hydrogel constructs (0.8 kPa, 600 mJ/cm²), the stiffness increased over time, reaching approximately 2.3 kPa after 30 minutes. This dual crosslinking strategy provides an optimal biofabrication approach by enabling spatial control of matrix structure through photocrosslinking, and subsequent enhancement of mechanical properties via secondary crosslinking.

### FLight printing using pristine protein-based photoresin

Next, we assessed the printability of pristine protein-based photoresins using FLight bioprinting. Given the crucial role of the photoinitiator in determining printing resolution and windows, we prepared a series of photoresins by varying the concentration of Ru and the ratio of Ru to SPS. Photorheological results revealed a positive correlation between Ru concentration and polymerization rate, with higher concentrations of photoinitiator (0.5 mM Ru and 5 mM SPS) leading to faster polymerization (**Figure S4**). Additionally, the ratio of Ru to SPS influenced the rate of photocrosslinking, with higher SPS concentrations at equivalent Ru levels accelerating photopolymerization.

Increasing photoinitiator concentrations results in faster photopolymerization, but printing resolution is also a critical factor in light-based biofabrication. To evaluate this, we printed the ETH Zurich logo and compared the feature sizes in the designed pattern to those in the final printed hydrogel, using a metric defined as the shape index (**Figure 1d**). The shape index, which measures printing fidelity, indicated that a Ru concentration of 0.1 mM provided better resolution (smallest positive feature size of hydrogel constructs), and a reduced SPS concentration resulted in improved printing fidelity (a shape index closer to 1.0). However, achieving this required higher light doses, which increase the potential for cell damage due to excessive light. Balancing these factors, we selected a 0.2 mM concentration of ruthenium and a 0.2 mM concentration of SPS as the optimized photoinitiator formulation for subsequent studies.

Before applying pristine protein-based photoresins in tissue engineering, it is essential to consider the potential cytotoxicity of the Ru-SPS photoinitiator. Previous studies have shown that Ru-SPS concentrations below 10 µM for 1 hour do not negatively affect cell viability or proliferation across various cell types. After FLight printing, ruthenium was removed by washing with PBS for 20 minutes (**Figure S5**), reducing its concentration to approximately 5 µM, as determined by optical absorbance compared to a standard curve. Overall, we demonstrated the potential of protein-based photoresins in FLight printing, enabling the creation of hydrogel structures with dimensions smaller than 150 µm, which is otherwise challenging to achieve with conventional crosslinking approaches such as gel casting.

### Microarchitectures and fibril structures formed in pristine protein-based FLight matrices

Compared to bulk hydrogels, the microarchitectures in FLight hydrogels provide efficient cell-guiding cues for the biofabrication of aligned tissues, such as skeletal muscle. To assess the presence of these microarchitectures when using protein-based photoresins, we used fluorescently labeled collagen and fibrinogen photoresins to print FLight matrices and performed confocal laser scanning microscopy. As controls, we also prepared fluorescently labeled GelNB-GelSH photoresins and bulk collagen hydrogels. As shown in **Figure 2a**, both bright-field microscopy and confocal laser scanning confirmed the formation of microarchitectures in FLight-printed hydrogel constructs. In contrast, no microarchitectures were observed in bulk hydrogel samples produced via gel casting.

**Figure 2.**
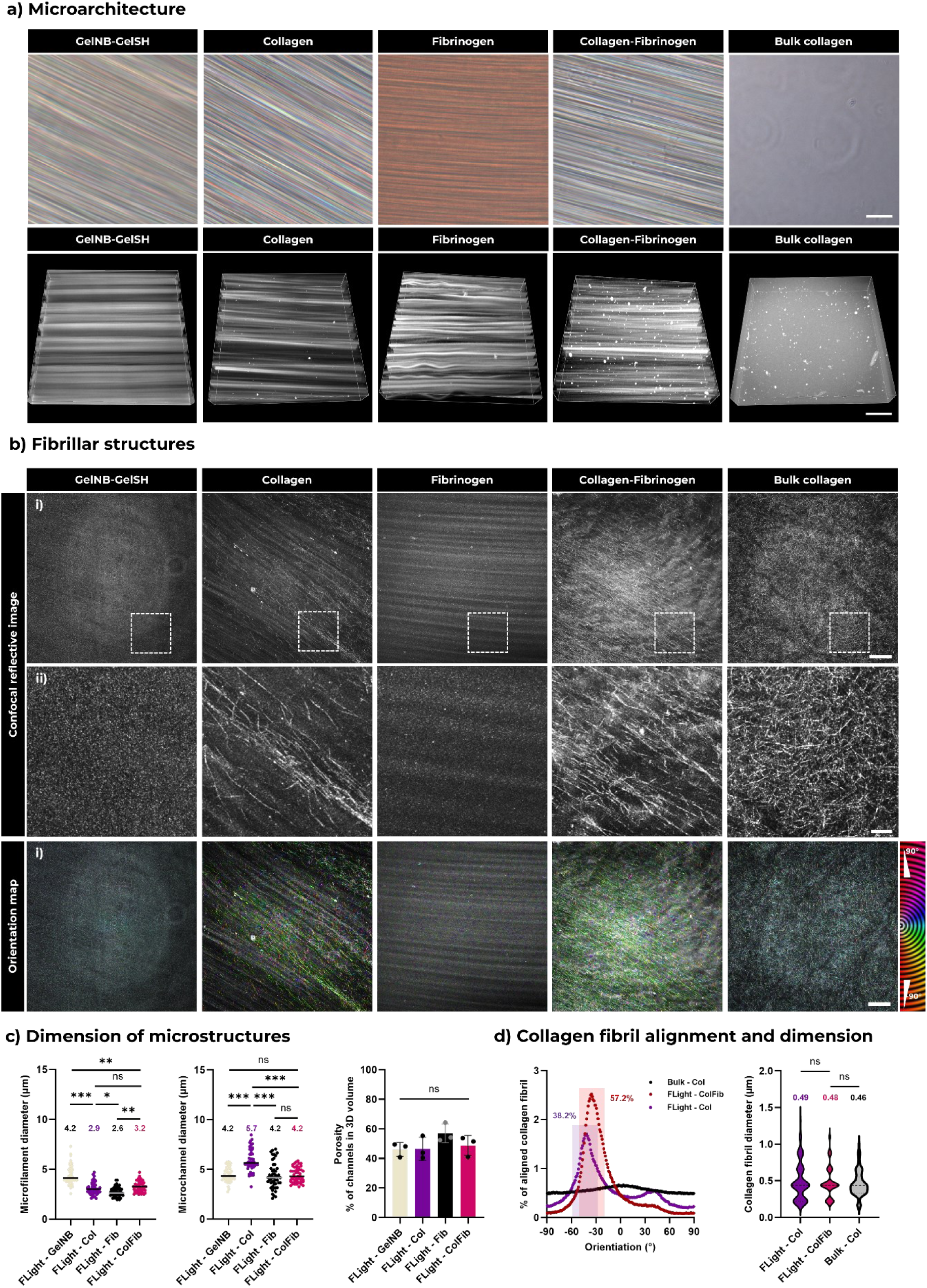
Microarchitectures and fibril structures formed in pristine protein-based FLight matrices. **a**) Bright-field and confocal laser scanning microscopy images of microarchitectures in different matrices. Scale bar: 50 µm. **b**) Reflective confocal microscopy images of fibril structures formed in different matrices after 60 min of secondary crosslinking at 37 °C. Orientation maps were generated using the Orientation J plugins in ImageJ, where the color indicates the orientation of the fibril structures. Scale bars: 5 µm in i), and 1 µm in ii) zoomed view. **c**) Characterization of microfilament diameters, microchannel diameters and the porosity of different matrices. Porosity was calculated from the volume ratio of microchannels to total hydrogel volume. All data were measured from the confocal scans following 3D reconstruction and segmentation (n=3, data size ≥50). **d**) Assessment of the fibril structure in different collagen matrices, including fibril orientation and the fibril diameter distribution. All data were collected from the reflective confocal scanning (for diameter distribution: n=3, data size: ≥50). The colored box highlights the aligned fibril along the direction of projection, where the alignment is defined as an angle between −10 and 10° from the peak of the orientation distribution. Abbreviations shown as: Col: collagen; Fib: fibrinogen; ColFib: collagen-fibrinogen. Data of microstructure dimension in c) are presented as mean ± SEM and individual points (*N* ≥ 3 with dataset size ≥ 50); \**P* < 0.05, \*\**P* < 0.01, \*\*\**P* < 0.001.

Analysis of the microarchitectures revealed that the microfilaments in ColFib hydrogel matrices were relatively smaller than those in GelNB-GelSH FLight matrices (**Figure 2c**). Overall, microfilament diameters in all matrices ranged from 2 to 8 µm, while the distribution of microchannel diameters ranged similarly from 2 to 10 µm. Specifically, the mean microfilament diameter was 4.2, 2.9, 2.6 and 3.2 µm in GelNB, collagen (Col), fibrinogen (Fib) and ColFib FLight matrices. The average diameter of microchannels was 4.2, 5.7, 4.2 and 4.2 µm in these matrices. Furthermore, no significant differences in porosity were found across all hydrogel matrices, with microchannels occupying approximately 50% of the total 3D volume of the hydrogel.

Given the previous observation of secondary thermo-crosslinking of collagen within the photocrosslinked hydrogels, we further investigated the formation of collagen fibril structures in FLight matrices. The fibril structures were examined with reflective confocal microscopy at a wavelength of 488 nm after secondary crosslinking.[31,32] Interestingly, we detected fibril structures in all hydrogels containing collagen; however, the fibrils in the bulk collagen hydrogel exhibited a random orientation (**Figure 2b**). In contrast, the collagen fibrils in the FLight-printed hydrogels were observed to be more aligned. Approximately 38% of the fibrils in the FLight-Col hydrogels were aligned within a range of –10° to 10° along the long axis of the microfilaments, while in the FLight-ColFib hydrogels, the aligned fibrils accounted for approximately 52% (**Figure 2d**). A possible reason for the increased collagen fibril alignment was that the microchannels in the ColFib matrix are narrower, which may confine fibril formation to the direction along the microfilaments. Furthermore, no significant differences were detected in the diameter of the fibril structures across the different collagen-based matrices, with a mean diameter of approximately 500 nm. This range is consistent with previous reports on the dimensions of *in vitro* collagen fibrils formed by self-assembly.[32]

### Pristine protein-based FLight matrix supports Pax7 expression and proliferation of Pax7-nGFP Primary myoblasts

After confirming the presence of microarchitectures, we encapsulated mouse Pax7-nGFP primary myoblasts in both pristine ColFib and functionalized GelNB FLight matrices to evaluate cell behavior (**Figure 3**). The selected gelatin photoresin formulation had a concentration similar to that of the pristine protein-based photoresin, approximately 2.8% w/v (GelNB: GelSH = 1: 1). In addition, the light dose (∼280 mJ/cm²) was optimized to ensure that the stiffness of the GelNB hydrogel matrices was comparable to that of the ColFib hydrogel (∼2.5 kPa). On day 0, before resuspending the cells in photoresin and encapsulating them in 3D FLight matrices, approximately 87% of the total cells expressed Pax7, as detected by GFP signal from the nucleus (MyoD^-^: 26%, MyoD^+^: 61%, see **Figure 3b**). Immunofluorescent staining for Ki67 indicated that more than 37% of the myoblasts were in an active proliferative state.

**Figure 3.**
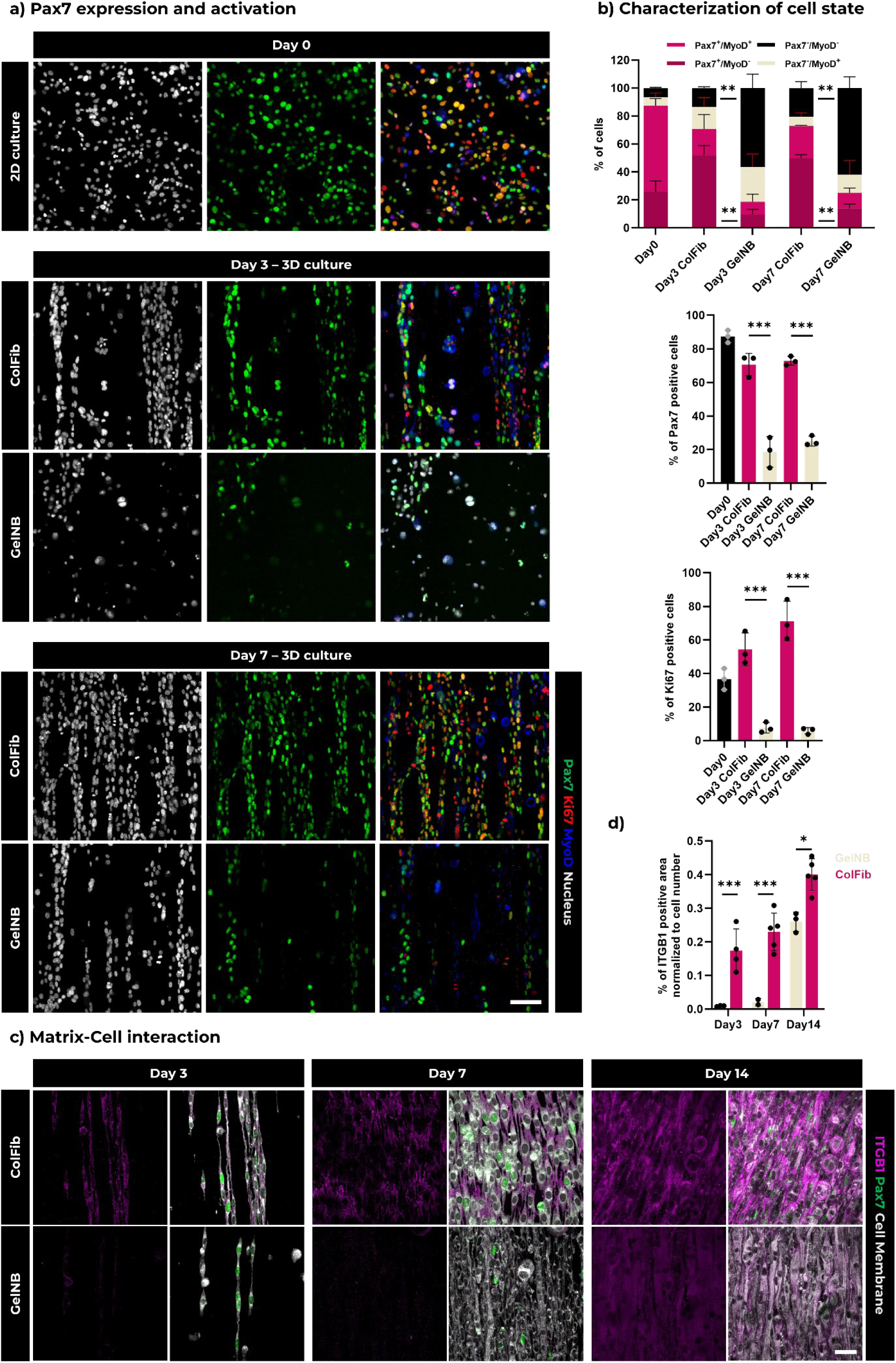
Pax7 expression and proliferation of Pax7-nGFP primary myoblasts is enhanced in ColFib FLight matrices compared to GelNB. **a**) Representative immunofluorescent staining of Pax7-nGFP primary myoblasts, when cultured on 2D surfaces (day 0), and encapsulated in pristine protein-based FLight matrices (ColFib) compared to gelatin matrices (GelNB-GelSH). Green: Pax7-nGFP; Red: Ki67; Dark blue: MyoD; and Gray: Nucleus. Scale bar: 100 µm, applicable to all images. **b**) Characterization of the state of Pax7-nGFP primary myoblasts in different FLight matrices at days 3 and 7 compared to 2D monolayer culture. All data were measured from the confocal scanning images in a) (n=3). **c**) Representative confocal images of Pax7-nGFP primary myoblasts encapsulated in different FLight matrices that show expression of integrin beta 1 (ITGB1) after 3, 7, and 14 days of culture. Pink: ITGB1; Green: Pax7-nGFP; Gray: Cell membrane (CellMask dye). Scale bar: 50 µm. **d**) ITGB1 positive area normalized to cell number in pristine protein-based FLight matrices (ColFib) vs gelatin matrices (GelNB-GelSH). All data were collected from the confocal scanning of c) (n≥3). Data in b) and d) are presented as mean ± SEM; \**P* < 0.05, \*\**P* < 0.01, \*\*\**P* < 0.001.

Notably, significant differences were observed between the ColFib and GelNB FLight matrices after 3 and 7 days of culture. In the ColFib matrix, the proportion of Pax7^+^ primary myoblasts remained relatively stable at 70% and 72% on day 3 and day 7, respectively (MyoD^-^: 18%, MyoD^+^: 52% on day 3; MyoD^-^: 22%, MyoD^+^: 50% on day 7). In contrast, the Pax7^+^ cell population in the GelNB matrix dropped to 19% and 24% on day 3 and 7, respectively. . Additionally, approximately 56% and 62% of the total cell population in the GelNB matrix expressed neither Pax7 nor MyoD (i.e., Pax7^-^/MyoD^-^), suggesting that cells in the GelNB matrix gradually lost their stemness (i.e., Pax7 expression) and activation ability (i.e., MyoD expression). Confocal images of Ki67 staining further indicated increased proliferation in the ColFib matrix. The proportion of Ki67^+^ myoblasts was 54% and 71% in the ColFib matrix, compared to only 8% and 7% in the GelNB matrix on days 3 and 7, respectively.

We hypothesized that the unmodified protein-based FLight matrices was better at promoting establishment of matrix-cell interactions, such as those involving the integrin family, compared to highly functionalized gelatin.[33] To investigate this further, we examined integrin beta 1 (ITGB1) expression in both matrices (**Figure 3c, d**). Immunofluorescent images showed that ITGB1 signals were more pronounced in ColFib matrices than in GelNB matrices at all time points. Although further investigation is needed to fully explain the differences in Pax7 expression and proliferation, the findings on ITGB1 suggest that, similar to *in vivo*, insufficient matrix-cell signaling may result in reduced Pax7 expression and proliferation.[6–8] In summary, while FLight bioprinting can successfully create microarchitectures that guide cell alignment, the chemical composition of the filamented matrices also appears to play a critical role in determining the cell state of Pax7-nGFP primary myoblasts.

### Maturation and functionalization of engineered FLight muscle constructs

Differentiation was initiated on day 7 by switching the growth medium (GM) to differentiation medium (DM) containing 2% horse serum. The maturation of the engineered FLight muscle was then assessed on days 11, 14, and 21. Immunofluorescence imaging of sarcomeric alpha-actinin (SAA) and F-actin revealed the formation and maturation of sarcomeres, the basic contractile units of muscle responsible for force generation (**Figure 4a**). Morphological analysis of these sarcomeric structures showed a more uniform distribution and length (close to 2 µm) over the 3 weeks of culture (**Figure 4c**). Additionally, myotube maturation and basal membrane-associated ECM deposition were evaluated by immunofluorescence staining using anti-myosin heavy chain and anti-laminin antibodies. We observed the fusion of myoblasts into myotubes with multiple myonuclei, accompanied by the formation of layered laminin structures (**Figure 4b**). In 3D reconstructions of confocal scans, Pax7^+^ nuclei were found adjacent to the myotubes and covered by laminin, resembling the satellite cell niche microenvironment observed *in vivo*.

**Figure 4.**
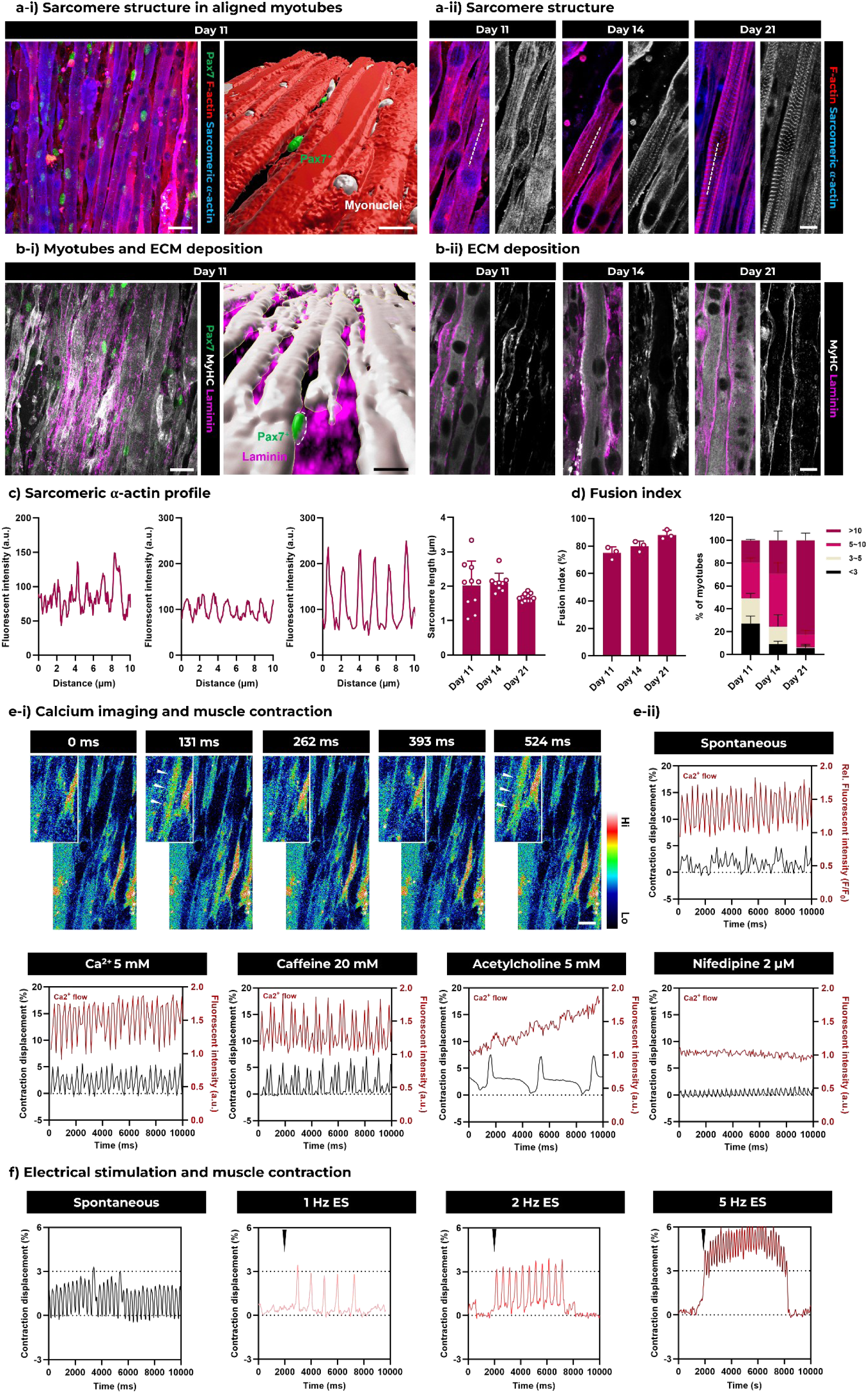
Maturation and contraction of engineered FLight muscle. **a-i**) Representative immunofluorescence images of engineered FLight muscle biofabricated using ColFib photoresin after 11 days of incubation (differentiation started at day 7). Green: Pax7-nGFP; Red: F-actin; Light blue: Sarcomeric alpha actinin (SAA). 3D reconstruction of the confocal scan showing the fused myotube, myonuclei and Pax7^+^nuclei in FLight hydrogels. Scale bar: 20 µm. **a-ii**) Representative immunofluorescence images of muscle fibers formed in FLight matrices. SAA distribution (in gray) highlights the maturation of sarcomeres at different time points. Scale bar: 10 µm. **b-i**) Representative immunofluorescence images showing laminin deposition of engineered FLight muscle biofabricated with ColFib photoresin after 11 days of incubation (differentiation started at day 7). Green: Pax7-nGFP; Gray: Myosin heavy chain; Pink: Laminin. And 3D reconstruction of confocal scanning showing the fused myotube, newly deposited laminin and Pax7^+^nuclei. Scale bar: 20 µm. **b-ii**) Representative immunofluorescence images of laminin deposition in engineered FLight muscle. Laminin distribution (in gray) highlights the deposition of basal membrane-associated ECM. Scale bar: 10 µm. **c**) Intensity distribution of SAA on the white dashed lines depicted in a-ii). From left (day 11) to right (day 21) indicates the more uniform distribution of SAA during the culture. **d**) Quantitative analysis of the fusion index of engineered FLight muscle after 11, 14, and 21 days of culture; and the percentage of myotubes with different numbers of nuclei (n=3). **e-i**) Representative fluorescence images of Ca^2+^ indicator at different time points (a higher fluorescence intensity upon Ca^2+^ binding). The color bar indicates the fluorescence intensity. Scale bar: 20 µm. **e-ii**) Fluorescence intensity of Ca^2+^ indicator (F/F0) and contraction displacement (%) of engineered FLight muscles under spontaneous contraction, with additional calcium chloride (Ca^2+^), caffeine, acetylcholine (ACh), and nifedipine added to the medium. **f**) Contraction analysis of engineered FLight muscles under spontaneous contraction, 3V-1Hz, 3V-2Hz and 3V-5Hz of electrical stimulation. The black arrows indicate the starting points of electrical stimulation.

Further assessment of the myoblast fusion index, defined as the ratio of fused myonuclei to the total number of nuclei, revealed a time-dependent increase in fusion. The fusion index increased from approximately 75% on day 11 to 90% after 21 days of culture (**Figure 4d**). Analysis of the number of fused nuclei per myotube showed a significant increase in the percentage of myotubes containing more than 10 myonuclei after 21 days in culture (increasing from 20% to 80%). These results collectively demonstrate the successful differentiation and fusion of Pax7-nGFP myoblasts in ColFib FLight matrices.

Interestingly, spontaneous muscle contractions were observed as early as day 14 (**Video S1**). To further investigate the contractility of the engineered FLight muscle, we examined muscle contractions and calcium transients. Calcium imaging during spontaneous contractions was first performed using a calcium indicator. In **Figure 4e**, the arrows indicate temporal changes in fluorescence intensity upon the binding of the indicator to calcium ions (Ca^2+^), confirming the presence of calcium transients (also **Video S2**). Although the spontaneous contractions exhibited irregular frequency and strain (∼3% of the total contraction displacement, ΔL/L x 100%), the contraction peaks closely overlapped with peak fluorescence intensity of the calcium indicator.

Based on these observations, we introduced several molecules related to muscle performance and calcium handling into the culture medium, including calcium chloride (Ca^2+^), caffeine, acetylcholine (ACh), and nifedipine (a commercial drug acts as calcium channel blocker) (**Video S3**).[34–37] The addition of extra Ca^2+^ and caffeine resulted in more uniform and pronounced contractions, characterized by more regular contraction patterns and larger displacement of muscle contraction (∼5%). Caffeine enhances muscle performance by increasing contraction speed and strength through the opening of Ca^2+^ channels in the sarcoplasmic reticulum, leading to the release of additional Ca^2+^.[38] Interestingly, while muscles exposed to caffeine exhibit an increased ability to recruit calcium, their relaxation and recovery speeds are diminished.[39] The addition of a high concentration of Ach led to even stronger contractions in the engineered FLight muscle, with peak contraction displacement reaching approximately 7%. In skeletal muscle, acetylcholine binds to nicotinic acetylcholine receptors (nAChRs), triggering their activation and causing the transience of cations (Na^+^ and Ca^2+^), which depolarizes the membrane of muscle fiber and initiates contraction. Acetylcholine then dissociates from the nAChRs and is degraded by acetylcholinesterase, preventing sustained nAChRs activation.[36] Interestingly, a plateau phase in the contraction curve was also observed, likely corresponding to depolarization induced by the sustained activation of the nAChRs by the high concentration of ACh.[36] In contrast, blocking calcium channels in nifedipine-treated engineered FLight muscle resulted in reduced calcium transients and a significant decrease in muscle contraction. Similar results have been observed *in vivo*: the application of nifedipine to rat muscle fibers reduces muscle contraction force, with the inhibitory effect of nifedipine on fiber contraction being positively correlated with its concentration.[40]

We then evaluated the contractile behavior of the engineered FLight muscle under electrical stimulation. A direct current (square wave) of 3V with varying frequencies was applied along the longitudinal axis of the muscle fibers. The results indicated that contraction frequency is modulated by the frequency of stimulation, with increased muscle contractions observed as the frequency increased (**see Figure 4f**). At a stimulation frequency of 5 Hz, tetanus-like contraction behavior was observed.

### Evaluation of the self-renewal potential of engineered FLight muscle after CTX-induced injury

One of the most important roles of satellite cells *in vivo* is to regenerate muscle after damage. We next used the mature FLight muscle model to explore the role of its Pax7^+^ cell population in muscle self-renewal following injury. Since regeneration relies on the Pax7^+^ cell population, we also fabricated GelNB-based FLight muscle for comparison, to explore differences in Pax7 states and regeneration outcomes. Muscle injury in the engineered FLight muscle was induced using cardiotoxin (CTX), a widely established *in vivo* and *in vitro* injury model. The cell state of the engineered FLight muscle was assessed before and after 2 hours of CTX treatment using immunofluorescence staining (**Figure 5a**). Prior to CTX treatment, a higher ratio of Pax7^+^ cells was observed in the total cell population within ColFib matrices (**Figure 5b and S6**) compared to GelNB matrices, consistent with previous findings of reduced Pax7 expression. Following CTX treatment, the percentage of Pax7^+^ cells in both matrices increased over time. However, a significant difference emerged after 7 days of recovery, with approximately 40% Pax7^+^ cells in the ColFib matrix compared to only 5% Pax7^+^ cells in the GelNB matrix. The proportion of Ki67^+^ cells within the total Pax7^+^ cell pool also increased over time, indicating the proliferation of Pax7^+^ cells.

**Figure 5.**
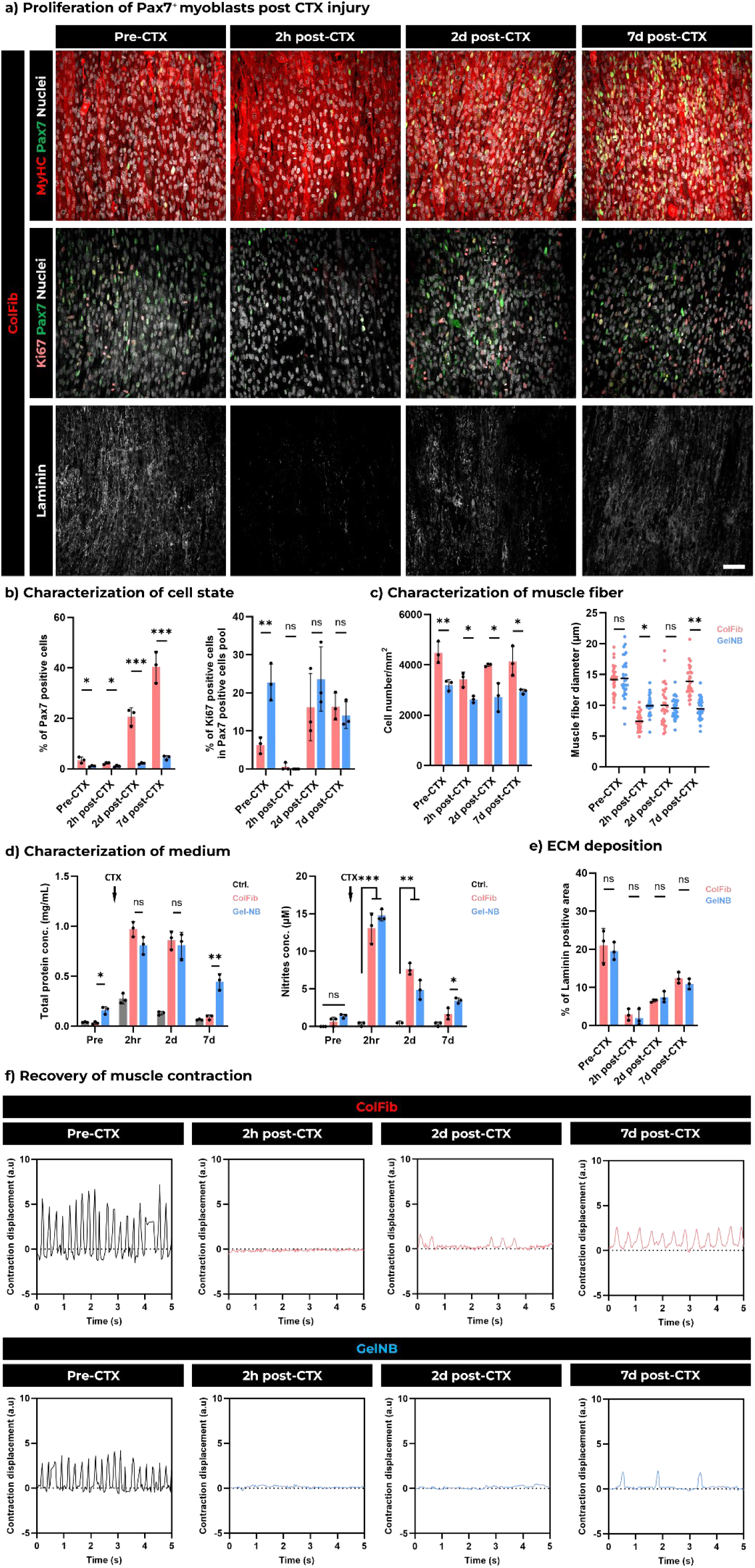
Self-renewal potential of engineered FLight muscle post CTX-induced injury. **a**) Representative confocal images of engineered FLight muscle after 21 days of culture (Pre-CTX), and 2 hours (2h post-CTX), 2 days (2d post-CTX) and 7 days (7d post-CTX) after CTX treatment. Scale bar: 50 µm. **b**) Quantitative analysis of Pax7^+^ cell population (% of Pax7^+^nuclei in the total nuclei) and Ki67^+^ cell population (% of Ki67^+^nuclei in the total nuclei) at different time points. All data were obtained from the 3D reconstruction of confocal laser scans (n=3). **c**) Characterization of cellularity and muscle fiber diameter. Cellularity was measured using ImageJ and muscle fiber diameter were calculated from 3D reconstruction of confocal images (MyHC signal, n=3, data size ≥20). **d**) Characterization of total protein concentration and nitrite concentration in the medium at different time points (n=3). **e**) Evaluation of laminin deposition before and after CTX-induced injury. All data were measured from immunofluorescence images using ImageJ (n=3). **f**) Spontaneous contraction analysis of engineered FLight muscles biofabricated from ColFib and GelNB at 2h, 2 days and 7 days after adding CTX. Data from b) to e) are presented as mean ± SEM; \**P* < 0.05, \*\**P* < 0.01, \*\*\**P* < 0.001.

Regarding the myotubes in the engineered FLight muscle, a decrease in cell density and muscle fiber diameter was observed immediately following treatment (2h post-CTX). However, a time-dependent recovery in cellularity and muscle fiber diameter was noted at days 2 and 7. Notably, the recovery of muscle fibers was more pronounced in the pristine protein matrix (**Figure 5c**), with significantly thicker muscle fibers in ColFib matrices (∼14 µm in diameter) compared to GelNB matrices (∼9.5 µm) on day 7. A significant reduction in laminin was also observed after CTX treatment, followed by a gradual increase on days 2 and 7, indicating that ECM deposition was initiated as the self-renewal of the engineered FLight muscle progressed (**Figure 5e**).

Additionally, total protein and reactive nitrogen species concentrations in the medium were measured. The concentration of both increased significantly immediately after CTX treatment, regardless of the matrix used (**Figure 5d**). Total protein likely included creatine kinase, and reactive nitrogen species, which are released from muscle upon injury and play a key role in activating satellite cells and the immune system to aid in muscle regeneration.[41–43] This may also explain the increased proliferation of Pax7^+^ cells. As the self-renewal of the engineered FLight muscle progressed, the concentrations of total proteins and nitrogen species gradually decreased on days 2 and 7.

Muscle contractility is another key indicator of functional recovery after injury. No spontaneous activity was detected 2 hours after the addition of CTX, but contractions were observed in the engineered FLight muscle biofabricated with ColFib photoresin on days 2 and 7 (**Figure 5f, Video S4**). Although the frequency and contraction displacement differed from pre-treatment levels (3% vs. 5%), these results indicate that muscle function was in the process of recovery. In contrast, for engineered FLight muscle created with GelNB, only minor and irregular contractions were observed on day 7 (contraction strain < 2.5%, also **Video S5**). The differences in the regeneration of muscle function were consistent with previous observations of muscle fiber diameter and may be attributed to variations in the Pax7^+^ cell pools.

In conclusion, the self-renewal potential of engineered FLight muscle was demonstrated by the presence of a Pax7^+^ cell population, muscle fiber morphologies, ECM remodeling, and most importantly, the recovery of muscle function, evidenced by contractility. The engineered FLight muscle holds significant potential as an *in vitro* muscle model.

### Enhanced self-renewal potential of engineered FLight muscle using small molecules

Finally, we explored the potential of engineered FLight muscle as an *in vitro* muscle model to evaluate the effects of small molecules on muscle regeneration. Previous studies on skeletal muscle regeneration using stem cell therapy have shown real potential for a small molecule cocktail (Forskolin, Chir 99021, and Repsox) in terms of clinical applicability and safety.[44] Therefore, we designed a muscle injury experiment to test the ability of these small molecules to improve muscle recovery after CTX injury. We extended the duration of CTX treatment to 4 hours to induce more severe muscle injury and investigated how low and high doses of Forskolin, Chir 99021 and Repsox affected Pax7^+^ cell pool and cell proliferation.

Pax7 and Ki67 expression after 7 days of treatment was assessed by immunofluorescence staining (**Figure 6a, c**). Compared to the untreated control group, the percentage of Pax7^+^ cells significantly increased in the engineered FLight muscles treated with low doses of Forskolin, Chir 99021, and both low and high doses of Repsox. The percentage of Ki67^+^ cells was significantly increased in the Forskolin (low/high) and Repsox (low/high)-treated groups compared to the control. However, reduced proliferation was observed in the Chir 99021 (high) group. These differences may be attributed to the distinct roles and functions of these molecules. For example, Forskolin is a cAMP pathway activator that enhances the proliferative potential of muscle stem cells.[45] Activation of cAMP signaling has been shown to slow the progression of muscle atrophy in dystrophic *mdx* mice by enhancing satellite cell proliferation.[46] While more detailed studies are needed to fully understand how cAMP signaling specifically aids muscle regeneration, it already represents a promising avenue for developing new therapeutic drugs.[46] Chir 99021 is a highly selective inhibitor of glycogen synthase kinase 3 (GSK3) and an activator of Wnt signaling, which can drive satellite cells from proliferation to differentiation by switching Notch signaling to canonical Wnt signaling.[47] In addition, activated satellite cells face a fate decision between self-renewal and differentiation.[48] Wnt signaling plays a dual role in this process: 1. it promotes the differentiation and myogenic function of muscle stem cells during regeneration; 2. It inhibits myogenic lineage progression, forcing satellite cells toward alternative lineages, which can lead to fibrosis in aging muscles.[49,50] Repsox, on the other hand, acts as a promoter of Notch signaling, which is crucial for maintaining Pax7 expression and the satellite cell pool.[51] *In vivo,* Notch signaling is activated during muscle injury, and failure to activate the Notch pathway can result in defective muscle regeneration.[48,52] A recent study has also shown that intramuscular injection of Notch 1 antibody can rescue regeneration defects in aged muscles.[53]

**Figure 6.**
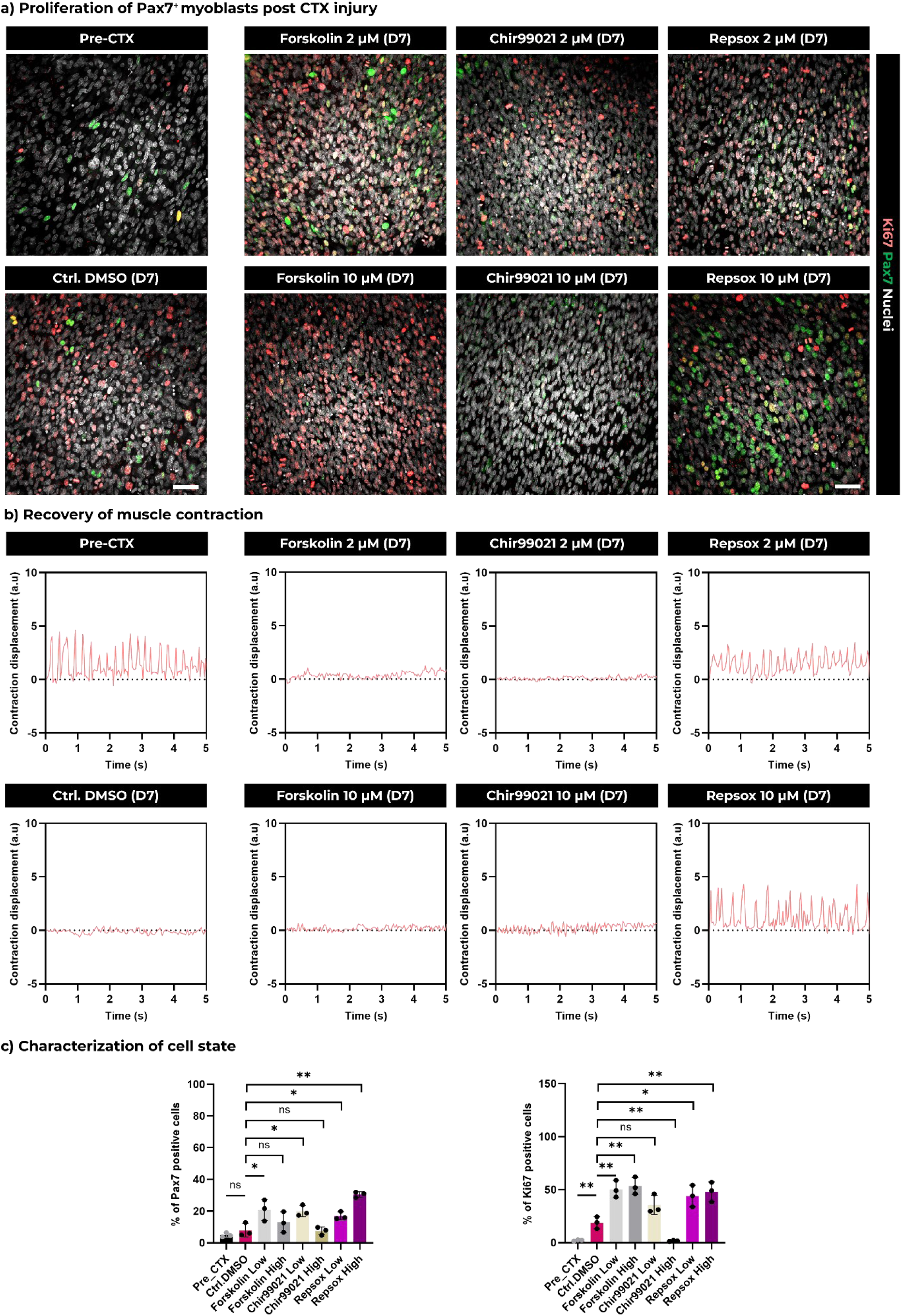
Enhanced self-renewal potential of engineered FLight muscle by different molecules. **a**) Representative confocal images of engineered FLight muscle before CTX treatment (Pre-CTX) and 7 days post CTX treatment with different small molecules and concentrations (Forskolin, Chir99021, Repsox). Scale bar: 50 µm. **b**) Spontaneous contraction analysis of engineered FLight muscles before and after 7 days of recovery under different culture conditions. **c**) Quantitative analysis of Pax7 positive cell population (% of Pax7^+^nuclei in the total nuclei) and Ki67 positive cell population (% of Ki67^+^nuclei in the total nuclei) under different culture conditions. All data were obtained from the 3D reconstruction of confocal laser scans (n=3). Data in c) are presented as mean ± SEM; \**P* < 0.05, \*\**P* < 0.01.

Furthermore, it has been reported that the addition of Forskolin, Chir 99021, and Repsox during the reprogramming of fibroblasts *in vitro* can lead to the upregulation of Notch pathway-related genes.[54] Given the pivotal role of the Notch pathway in regulating Pax7 expression, we performed qRT-PCR to evaluate the expression of Notch pathway and muscle regeneration-related genes (**Figure S7-8**). The results indicated a correlation between the expressions of Notch1/3, Hey1, and Pax7. Specifically, the lowest expression of Pax7-related genes (Notch1/3, Hey1) and Ki67 was detected in the Chir 99021 (high)-treated engineered muscle, consistent with the trends observed in the immunofluorescence staining studies. Additionally, the expression of genes associated with muscle maturation and contraction (Myh1, ATP2A1, PPARGC1A) was increased only in the engineered muscle treated with Repsox at both low and high concentrations. Notably, only the engineered FLight muscle treated with Repsox (low/high) showed a recovery of spontaneous contractions (**Figure 6b, Video S6**).

Overall, at both the gene and protein levels, we observed that different molecules led to different regeneration outcomes, underscoring the need for further investigation into these drugs and their underlying mechanisms. Nevertheless, the engineered FLight muscle demonstrated significant potential as a drug screening platform, advancing the development of new therapies for skeletal muscle diseases.

## Conclusion and Discussion

In this study, we engineered *in vitro* muscle constructs using pristine collagen-fibrinogen photoresin and Pax7-nGFP primary myoblasts. Compared to chemically modified gelatin-based photoresin, the Pax7^+^ cell pool was better maintained within the ColFib FLight matrix. After three weeks of maturation, we observed aligned, contractile muscle fibers resembling *in vivo* organization, surrounded by Pax7^+^ cells. The observation of spontaneous contraction and responses to electrical stimulation demonstrated the function of these engineered *in vitro* muscles. In addition, the self-renewing property of engineered muscles following chemically induced injury was shown to be dependent on the involvement of Pax7^+^ cells. This repair process could be further enhanced by small molecules, such as Notch pathway promoters. This engineered *in vitro* muscle model holds significant promise as a high-throughput platform for advancing our understanding of muscle physiology and pathology.

Future work could focus on further development of the bioactive photoresin by incorporating additional components such as laminin, fibronectin, or skeletal decellularized extracellular matrix (dECM). This direction holds promise for creating an even more biomimetic and *in vivo*-like matrix environment, which could better preserve the stem cell niche. Recent studies have identified new signaling pathways activated in the stem cell niche, such as integrin alpha 7, a laminin-mediated integrin subtype that regulates the symmetric and asymmetric division of satellite cells.[55] Proper regulation of these two types of division could support long-term differentiation while maintaining muscle stem cell quiescence *in vitro*. Conversely, selectively disrupting these matrix-cell interactions (e.g., through integrin-blocking antibodies or gene editing) could enable the biofabrication of disease-specific *in vitro* models for certain muscle disorders.[10,56]

In addition to changing the biomaterials, optimization of microstructures and their dimensions may offer a flexible matrix platform for various applications. For example, creating larger microchannels could support the formation of thicker muscle fibers, particularly when the biofabrication strategy is applied to human muscle tissue, which typically exhibits thicker muscle fibers than those in mice, with diameters ranging from 50 to 100 µm.[57] In our previous studies, using photomasks and customized projection patterns from a digital micromirror device (DMD) system, we successfully generated larger channel structures. On the other hand, considering that the native stem cell niche presents a highly confined microenvironment, it remains to be investigated whether tuning microchannel diameters could alter the mechanical confinement applied to cells, potentially triggering specific mechanotransduction pathways and ultimately affecting Pax7 expression and stem cell quiescence. It has been reported that applying static compressive force to satellite cells leads to nuclear deformation and creates a quiescent muscle stem cell niche.[58]

From an engineering perspective, combining the engineered muscle model with other technologies could further expand its applications. For instance, performing the printing process directly on existing mechanical testing platforms could allow for a more comprehensive evaluation of *in vitro* muscle performance (e.g., contraction force), which is currently lacking in this work. Furthermore, considering the rapid development of non-invasive real-time monitoring technologies for *in vitro* systems, integrating biofabrication, tissue maturation, stimulation, imaging, and signal detection (e.g., electrochemical sensors of key protein-creatine kinase)[59] into a single engineered chip would significantly accelerate the development of new therapies for skeletal muscle diseases.

Development of engineered muscle tissues has been a highly prolific area of research, with significant implications for advancing muscle disease studies and drug development. Mini-muscle constructs, particularly those composed of myogenic progenitors derived from induced pluripotent stem cells (iPSCs), offer a promising avenue for creating patient-specific and physiologically relevant platforms to study disease mechanisms and test therapeutic interventions. Notably, iPSC-derived Pax7^+^ cells from Duchenne muscular dystrophy (DMD) patients, which lack dystrophin expression, allow for the biofabrication of muscle tissues that accurately recapitulate the disease phenotype of DMD.[60,61] In addition to DMD, these engineered muscle tissues offer new pathways for exploring various muscle pathologies, including myotonic dystrophy,[62] sarcopenia,[63] inflammatory and metabolic myopathies,[64] muscle fibrosis.[65]

In addition to drug screening, these platforms facilitate mechanistic studies to dissect the cellular and molecular processes underlying muscle biology and pathology. *In vitro* muscle models can be employed to investigate signaling pathways involved in muscle growth and atrophy, providing critical insights into potential therapeutic targets.[66] For example, muscle atrophy can be induced in cultured cells through disuse or denervation, enabling researchers to examine the molecular and cellular alterations associated with muscle wasting.[67] This approach can drive the discovery of novel strategies to combat atrophy. Moreover, these models can be used to study the development and function of different types of muscle fiber (e.g., fast and slow twitch), offering a deeper understanding of factors influencing fiber-type composition and their role in muscle diseases or adaptations.[68]

While this work primarily focuses on muscle tissues, the methodologies presented here can also be extended to the interfaces between muscle and other tissues, such as the neuromuscular or myotendinous junctions.[69,70] Embracing a multi-pathology modeling approach allows for the exploration of combinatorial therapies, where drugs targeting different aspects of muscle disease can be combined to achieve synergistic effects and improved patient outcomes. This approach aligns with the growing trend of precision medicine, where treatments are tailored to individual patients based on the genetic and molecular profiles of their conditions.

## Materials and Methods

### Synthesis of GelNB and GelSH

The synthesis protocol of GelNB and GelSH was adapted from previous work.[22] In brief, for the synthesis of GelNB, 25 g of gelatin type A from porcine skin (Sigma, G2500) was dissolved in a 0.3 M carbonate-bicarbonate (CB) buffer at 40 °C to prepare a 10 % w/v gelatin solution. Once the gelatin had fully dissolved, 0.5 g of cis-5-norbornene-endo-2,3-dicarboxylic anhydride (CA) crystalline powder was added to the reaction mixture, which was then stirred vigorously. The CA crystalline powder was added in a sequential manner, with a total amount of 2.0 g. The reaction solution was diluted two-fold in pre-warmed milliQ H_2_O, and the pH was adjusted to 7 before filtration (0.2 µm pore size). The solution was dialyzed (3.5 kDa cutoff cellulose tubing) for 3-4 days against milliQ H_2_O at 40 °C with frequent water changes before lyophilization. The degree of functionalization (DoF) was calculated using ^1^H NMR (Bruker Ultrashield 400 MHz, 1024 scans), and found to be about 0.091 mmol/g.

For GelSH synthesis, 25 g of gelatin type A was first dissolved in 1.25 L of 150 mM MES buffer (pH = 4.0) warmed up to 40 °C for a final concentration of 2 % w/v. When completely dissolved, 2.38 g (10 mmol) of 3,3′-dithiobis(proprionohydrazide) (DTPHY) was added to the reaction solution under stirring. Then, 3.83 g (20 mmol) of 1-ethyl-3-(3′-dimethylaminopropyl)carbodiimide hydrochloride (EDC) was added, and the reaction was left to proceed overnight at 40 °C. Tris(2-carboxyethyl)phosphine (8.6 g, 30 mmol) was then added to the reaction mixture, and the reaction was left to proceed for 12 h in a sealed flask under gentle stirring. The solution was filtered (0.2 µm) and dialyzed (3.5 kDa cutoff cellulose tubing) for 3-4 days against milliQ H_2_O balanced to pH 4.5 with diluted HCl at 40 °C before lyophilization. Gel-SH was stored under an inert atmosphere at −20 °C prior to use. The degree of functionalization (DoF) was determined by ^1^H NMR (Bruker Ultrashield 400 MHz, 1024 scans), and found to be about 0.774 mmol/g.

To synthesize Gelatin-Rhodamine (Gel-Rho, a fluorescently labeled derivative of GelNB), 1.0 g of GelNB was dissolved in 100 mL of 0.3 M sodium bicarbonate (pH = 9.0) at 40 °C overnight. 10.0 mg of rhodamine B isothiocyanate was dissolved in 1.0 mL dimethyl sulfoxide (DMSO) and added to the Gel-NB solution. The reaction was allowed to proceed at room temperature overnight. The product was purified by dialysis (3.5 kDa cut-off) against milliQ H_2_O at 40 °C for 3 days with frequent water changes, protected from light, and subsequently lyophilized.

### Preparation of GelNB photoresin

In order to prepare GelNB-based photoresin, the lyophilized GelNB and GelSH were dissolved in PBS solution at 37 °C with 1:1 molar ratio of NB to SH. Lithium phenyl-2,4,6-trimethylbenzoylphosphinate (LAP) was added as photoinitiator from a stock solution of 5 % w/v in PBS to obtain a final concentration of 0.05 % w/v. Photoresins were prepared with a total polymer concentration of 2.8 % w/v and filtered through 0.2 µm filters to remove scattering particles and sterilize photoresin prior to use.

### Preparation of pristine protein-based photoresin

In all experiments, lyophilized pure collagen type I (Advanced Biomatrix, 5005) was dissolved in CFI solution (ETH Zurich Invention Disclosure 2022-010, in progress) and fibrinogen (Sigma, F8630) was dissolved in 1× PBS solution unless otherwise stated. Collagen solutions were prepared at concentrations of 1-6 mg/mL, depending on the experiment. Fibrinogen solutions were prepared at concentrations of 10-50 mg/mL. To prepare the collagen-fibrinogen solution, a total protein concentration of 28 mg/mL (2.8% w/v) was prepared by mixing 6 mg/mL of the collagen solution with 50 mg/mL of the fibrinogen solution in a 1:1 mixture. To prepare photoresin, additional ruthenium and sodium persulfate (Ru-SPS, Advanced Biomatrix, 5248) was added to the protein solution to a final concentration of 0.2 mM photoinitiator (Ru:SPS = 1:1).

### Circular dichroism (CD) measurements

Circular dichroism (CD) spectra of pristine collagen type I dissolved in acetic acid, PBS and CFI solution were recorded using a Jasco J-815 CD spectrometer. Spectra were collected in the wavelength range of 190–260 nm. Collagen was prepared at a concentration of 0.1 mg/mL in the respective solutions and loaded into a 1 mm path-length quartz cuvette (Hellma). Measurements were performed at 4°C in continuous scanning mode with a bandwidth of 1 nm (n=3 for each condition). Buffer-baseline spectra were subtracted from the final measurements.

### Photorheology and Thermorheology

Photorheology analyses were carried out on an Anton Paar MCR 302e equipped with a 20 mm parallel plate geometry, 6 mm glass substrate, and OmniCure Series1000 lamp (Lumen Dynamics) used at 60% output power (equal to 2 mW/cm^2^) with narrow 405 nm bandpass filters (Thorlabs). To prevent the sample from drying during the measurement, a wet tissue paper was placed within the chamber. 80 µL of photoresin samples were loaded and left to equilibrate for 3 min prior to starting the analysis. Oscillatory measurements were conducted at a shear rate of 2% and a frequency of 1 Hz, with a gap of 200 µm and 10 s acquisition interval.

The temperature-dependent rheological properties of collagen and collagen-fibrinogen photoresin were evaluated using an Anton Paar MCR 302e rheometer, equipped with 20 mm parallel plate geometry and a stainless-steel floor. To prepare the collagen in different solutions, the following procedure was followed: 1. To prepare the collagen in acid (pH = 3.2), lyophilized collagen was dissolved in 20 mmol acetic acid to a concentration of 3 mg/mL. 2. Collagen in PBS: Lyophilized collagen was dissolved in a mixture of milliQ water and 10× PBS (9:1, v/v) at 4 °C to prepare a neutralized collagen solution with a final concentration of 3 mg/mL. 3. Collagen in CFI: The lyophilized collagen was dissolved in CFI solution (a glucose-phosphate-rich solution modified from previous work) to reach a final concentration of 3 mg/mL. For the rheological measurements, 80 µL of the solution was applied to the steel floor, the gap distance was set to 0.2 mm, and shear measurements were performed at a shear rate of 2% and a frequency of 1 Hz, with acquisition intervals of 10 seconds. During the initial 5 minutes of the measurements, the temperature was maintained at 4 °C. Thereafter, it was increased in 3 steps by the program to reach 37 °C (4 °C > 15 °C > 25 °C > 37 °C).

### Compression Modulus Measurements

To perform compressive tests, cylindrical hydrogel constructs were prepared using the FLight printer with a circular projection pattern (an array of 4 circles), each with a diameter of 5 mm and a height of 4 mm (along the microfilament direction). Uniaxial unconfined compression tests were performed on a Texture Analyzer (Stable Micro Systems, TA.XTplus), equipped with a 500-g load cell and a flat plate probe with a diameter of 15 mm. To ensure complete contact between the hydrogel samples and the plates, a preload of 0.2 g was applied prior to testing. Samples were compressed to a final strain of 15% at a strain rate of 0.01 mm/s. The elastic compressive modulus was calculated by linear fitting of the initial linear region (0.3-3%) of the stress-strain curve. All aforementioned tests were conducted at room temperature.

### UV-Vis Spectroscopy

Removal of ruthenium from printed FLight constructs was analyzed by the absorbance of ruthenium from PBS solution after 0, 5, 10, 20 min of washing. The optical absorbance of ruthenium was measured using a microplate reader (BioTek Synergy H1 Multimode Reader) from 300 to 700 nm with a scan bandwidth of 2 nm. The absorbance curves of the washing solution were compared with a standard absorbance curve of ruthenium in PBS solution at a concentration of 5 mM and 5 µM.

### Cell and Tissue culture

The mouse Pax7-nGFP primary myoblasts and associated culture protocols were kindly provided by Prof. Ori Bar-Nur. To culture and expand myoblasts in 2D environments, the culture flasks were first coated. In brief, the coating solution was prepared by diluting 1 mL of Matrigel (Corning, 356237) in 24 mL of low-glucose DMEM (Thermo Fisher Scientific, 31885023) with an additional 1% v/v Penicillin-Streptomycin (Thermo Fisher Scientific, 15140122). This solution was then stored at −20 °C for subsequent use. The coating solution was transferred to the flasks and shaken to ensure complete coverage of the surfaces. The flasks were then placed on ice for 5 minutes prior to a 1-hour incubation at 37 °C.

To culture and expand myoblasts, a culturing medium (CM) was prepared by first mixing Dulbecco’s Modified Eagle Medium (DMEM, Thermo Fisher Scientific, 41966029) with F-10 Medium (Thermo Fisher Scientific, 22390025) in a 1:1 ratio. The mixture was supplemented with 10% v/v horse serum (Thermofisher 16050122), 20% v/v fetal bovine serum (Gibco, 10270106), 1% v/v Penicillin-Streptomycin (Thermo Fisher Scientific, 15140122), and recombinant FGF2 at a concentration of 10 ng/mL (Bio-Techne, 233-FB-500). To differentiate primary myoblasts, a differentiation medium (DM) was used by adding 1% v/v Penicillin-Streptomycin (Thermo Fisher Scientific, 15140122), 1% v/v Insulin-Transferrin-Selenium (Corning, 25800CR) and 2% v/v horse serum (Thermo Fisher Scientific, 16050122) in DMEM (Thermo Fisher Scientific, 41966029).

The printed hydrogel constructs were initially cultured in CM for 7 days with medium changed every 2 days. For Pax7-nGFP myoblast differentiation, all cell-laden hydrogel samples were incubated in DM for 14 days. The DM was exchanged every 2 days for the week of differentiation and replaced with fresh medium everyday once spontaneous muscle contractions were observed. Without specific elaboration, all incubations were performed at 37 °C and a 5% concentration of CO_2_.

All added molecules were as follows: caffeine (Thermo Fisher Scientific, A10431.30), acetylcholine chloride (Thermo Fisher Scientific, L02168.14), nifedipine (Thermo Fisher Scientific, 329270010), forskolin (Sigma-Aldrich, F6886), CHIR 99021 (Sigma-Aldrich, SML1046), RepSox (Sigma-Aldrich, R0158). Caffeine, acetylcholine and nifedipine were diluted in culture medium. Forskolin, CHIR 99021 and RepSox were initially prepared according to the supplier’s protocol and then diluted in culture medium.

### Preparation of Bioresin

As previously indicated, a variety of photoresins were first prepared. The GelNB photoresin and Fibrinogen photoresin were filtered with a 0.2 µm filter (SARSTEDT AG, Filtropur S 0.2) in order to sterilize the photoresin and remove scattering particles. The filtration of the collagen solution was omitted for the preparation of Col-Fib photoresin due to its high viscosity. Subsequently, the collagen solution was mixed with the fibrinogen photoresin (in a 1:1 ratio), after which the Pax7-nGFP myoblasts were resuspended in the photoresin at a concentration of 1 million cells/mL.

### FLight Biofabrication

Cell-laden bioresin was first loaded into cuvettes (Thorlabs, CV10Q14F) or 8 well chamber slide (ibidi, 80801). The cuvettes and chamber slide were then transferred to 4 °C for 15 min (only applicable to the GelNB-based bioresin). The FLight printing process was conducted using a customized FLight3D printer (Lasertack 500 mW 405 nm Diode Laser, and Raspberry Pi with integrated control software), with a laser intensity of 55 mW/cm^2^. The projection pattern was designed accordingly, such as the ETH Zurich logo using Affinity Photo software with a resolution of 1920 x 1080 pixels (in PNG format, 8-bit grayscale). The projection pattern was then transferred to the FLight3D printer and loaded onto the built-in DMD controller. The light doses for FLight projection were optimized depending on biomaterials in order to match the matrix properties such as stiffness between different matrices. Immediately following the FLight printing, the cuvettes were warmed up in a heating bath at 37 °C (this step was only necessary for GelNB-based bioresin). The uncrosslinked photoresin was then removed via 3 times washing in pre-warmed PBS. To facilitate enzymatic crosslinking of Col-Fib hydrogel constructs, samples were incubated in a crosslinking solution comprising sterilized PBS and thrombin from bovine plasma (Sigma, T4648) at a final concentration of 5 units/mL for 30 minutes at 37 °C.

### Immunofluorescence Staining

FLight muscle constructs were washed with pre-warmed PBS solution three times and then fixed in 4% paraformaldehyde (PFA) for 1 hour at 4 °C. Subsequently, the engineered muscle samples were treated with 0.2% v/v Triton-X100 in PBS for 30 min (permeabilization), prior to being blocked with 1% w/v bovine serum albumin (BSA) in PBS for 2 h at 4 °C. These samples were then incubated with primary antibodies diluted in 1% w/v BSA-PBS for 12 h at 4 °C. Next, the samples were washed three times with PBS, incubated with diluted secondary antibodies, 1:1000 diluted Hoechst 33342, and phalloidin-tetramethylrhodamine B isothiocyanate working solution (0.13 µg/mL, Sigma, P1951) in 1% w/v BSA-PBS for 2 h at 4 °C (if applicable). More details on primary and secondary antibodies are listed in **Table S1**.

### Confocal Fluorescent Imaging

All confocal fluorescent imaging of hydrogel microstructures and (immuno)fluorescent imaging were performed on a confocal laser scanning microscope (Olympus, Fluoview FV3000). The microscope was equipped with 20× 0.75 NA (air) / 30× 1.05 NA (silicon) / 60×1.3 NA (silicon) objective lenses and detectors (4 GaAsP photomultiplier modules; 405 nm, 488 nm, 561 nm, 640 nm). Z-stack scanning was acquired from at least 20 µm depth of constructs at 2-5 µm step sizes. All images were captured using multiple excitation wavelengths and the emission signals were captured using suitable detectors. For example, the 405-nm detector was used to image nucleus, the 488-nm detector was utilized for imaging of GFP signal.

### Reflective Confocal Imaging and Analysis

Reflective confocal imaging was conducted on a Zeiss LSM 780 upright confocal microscope equipped with a 63× 1.4 NA (oil) objective (Zeiss, Plan-Apochromat DIC M27) and an argon laser (wavelength of 488 nm). To detect fibril structures from the deeper region of the hydrogel, all images were acquired at a depth of at least 100 µm from the surface of the hydrogel using Zeiss Zen microscopy software (resolution ≥ 1024 × 1024 pixels). For image analysis of collagen fibril structures, the raw image files were first loaded into the Zeiss Zen Blue analysis software, and a single stack image of the fibril was extracted and finally exported to ImageJ for subsequent analysis with fitting to a Gaussian distribution.

### 3D Reconstruction of Confocal Images

To conduct 3D reconstruction of confocal scans, Z-stack confocal images were initially converted to Imaris-compatible files using the Imaris File Converter software (Oxford Instruments, Ver. 10.0.0), which produces an .ims file. The Surface function was performed using Imaris (Oxford Instruments, Ver. 10.0.0) to create separate 3D reconstructed surfaces for different fluorescent signals (e.g., Hoechst 33342, GFP, Alexa 568 and Alexa 647). The surface resolution and threshold were set by default in the first 3D reconstruction process, without the application of a filter function. This configuration was then stored and applied to all subsequent 3D reconstructions.

2D plots of the fluorescent signal volume (on the x-axis) and the volume ratio of the fluorescent signal to the total 3D scanning volume (on the y-axis) were generated from Vantage function. To determine the ratio of microchannels in the total 3D hydrogel volume (i.e., porosity of hydrogel matrix), Z-stack confocal images of fluorescent-labeled microfilaments were reconstructed using the aforementioned method (n=3). The percentage of microchannels (*P*) was calculated using the following formula:

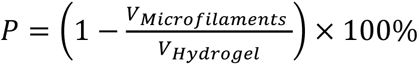

The total fluorescence positive area (*A_ITGB1_*) of integrin Beta I in 3D FLight hydrogel was measured by Imaris and normalized to the total number of cells (n=3).

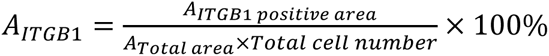

Similarly, the ratio of laminin positive area post CTX treatment was measured by Imaris (n=3).

### Analysis of microstructures

To characterize the microstructure, including the diameter of microfilaments and microchannels, the Z-stack confocal scans were first transformed into maximum-intensity projection images using the Z-project function in the Fiji software (https://fiji.sc/). To segment microfilaments, the default threshold was applied to all scans. A line perpendicular to the direction of the long axis of the microfilaments was drawn using the Straight function. The diameters of both microfilaments and microchannels were measured from the data presented in the Plot Profile panel of the Fiji software (n=3, dataset size ≥ 50). Similarly, the diameter of fibril structures was measured from the exported reflective confocal images using the Plot Profile function.

To assess the alignment of fibril structures formed in the FLight matrix, the reflective confocal stacks were first processed using maximum-intensity projections and the segmentation function as previously outlined. Then the OrientationJ plugin (available at http://bigwww.epfl.ch/demo/orientationj) was used to analyze the orientation distribution of the fibril structures. The structure tensor of the local window was set to 2 pixels and configured with a Gaussian gradient.

### Calcium Imaging

Intracellular calcium transients were assessed by incubating FLight muscle constructs with Rhod-2 AM dye (cell-permeable fluorescent Ca2^+^ indicator). Briefly, 10 µL of Rhod-2 AM calcium dye (Stock solution: 1 mg/mL, Abcam, AB142780) in dimethyl sulfoxide (DMSO) was carefully mixed with 10 µL of Pluronic F-127 (diluted in DMSO as 20% v/v). The resulting solution was added to 980 µL of DMEM (serum-free) to prepare imaging solution. Muscle constructs were washed with pre-warmed PBS and incubated with the imaging solution at 37 °C for 1 hour. Subsequently, the samples were washed with PBS three times and transferred to DM for calcium imaging.

To record the time-lapse calcium transients, the imaging was performed on a confocal laser scanning microscope (Olympus, Fluoview FV3000) using a resonant scanner with a fixed scanning resolution of 512 × 512 pixels. Samples were placed in an incubator chamber within the microscope to maintain a physiological condition (37 °C, 5% CO_2_). Each recording of muscle contraction and calcium flux exceeded 200 frames (approximately 130 ms per frame) and was initiated within 10 seconds after adding drugs/reagents (e.g., caffeine and nifedipine).

### Muscle Contraction and Analysis

Custom software was developed in Python to analyze muscle contraction, integrating a graphical user interface (GUI) using Tkinter (https://docs.python.org/3/library/tkinter.html). The software allows the user to choose a video file (e.g., from time-lapse microscopy images) in which the spatial coordinates of specific points will be tracked over time. The points can either be chosen manually by the user, or a selection of the best trackable points is presented to the user in the automatic mode. In automatic mode, points are selected using the Shi-Tomasi corner detection algorithm (https://docs.opencv.org/4.x/d4/d8c/tutorial_py_shi_tomasi.html), which is an improvement on the Harris Corner Detector and ensures optimal tracking points. The point coordinates were analyzed using the Lucas-Kanade optical flow method (https://docs.opencv.org/4.x/db/d7f/tutorial_js_lucas_kanade.html). All image analysis methods were implemented using the OpenCV library. The results were saved as time series data in CSV format using Pandas and visualized using MatPlotLib.

### Cardiotoxin (CTX) Injury

The matured muscle tissue constructs were washed with a pre-warmed PBS solution and then incubated in DM containing 2 µM CTX (Latoxan, L8102) for a period of 2 hours at 37 °C. To evaluate the effect of small molecules on the regeneration of muscle constructs, the engineered muscle tissues were treated with 4 μM CTX in DM for 4 hours. Following the CTX incubation, all tissue constructs were initially rinsed three times with pre-warmed PBS to remove any residual CTX. Subsequently, the tissue samples were transferred to fresh CM for 7 days of culture. At each of the time points of 2 hours, 2 days, and 7 days after the muscle tissues were placed in fresh CM, 500 μL of the CM was collected for analysis of total protein and reactive nitrogen species.

### RNA Isolation and Quantitative Reverse Transcription Polymerase Chain Reaction

The RNA isolation protocol of hydrogel constructs was adapted from previous work.[71] Briefly, FLight muscle constructs at different time points were transferred to 1.5 mL Eppendorf tubes and snap frozen using liquid nitrogen, then stored at −80 °C before RNA isolation. The RNA isolation was conducted by first adding 500 µL of NucleoZOL (Macherey-nagel, 740404.200) to tissue-laden tubes. The FLight muscle tissues were homogenized and then incubated with NucleoZOL on ice for 10 min. DNase-free water was added to the mixture, then centrifugated for 10 min at 12000 rcf. The supernatants were mixed with membrane-binding solution and isopropanol and then purified using ReliaPrep RNA Clean-up kit (Promega, Z1073). The yield and purity of the isolated RNA were determined with a spectrophotometer (Thermo Scientific, NanoDrop One). An A260/280 ratio of between 1.9-2.1 was accepted as adequate quality for the RNA samples. The isolated RNA was transcribed to complementary DNA following the instruction of GoScript Reverse Transcriptase kit (Promega, A5003). The relative gene expression levels were determined on real-time PCR System (Applied Biosystems, QuantStudio 3) with the GoTaq qPCR kit (Promega, A6001). The GAPDH housekeeper gene was used as an internal control for the normalization of RNA levels. Primer pairs used are listed in **Table S2**.

### Total Protein and Reactive Nitrogen Species Analysis

Total protein release from engineered muscle was assessed using Pierce 660nm Protein Assay Reagent (Thermo Fisher Scientific, 22660) according to the protocol provided by the manufacturer. Briefly, BSA protein standards and media samples were prepared by combining 10 µL of BSA/sample solutions with 150 µL of the reagent and loaded into a 96-well plate. These solutions were then gently mixed and incubated at room temperature for 5 minutes. Absorbance was measured at 660 nm on a plate reader (BioTek Instruments, Synergy H1). The total protein concentration of the samples was determined from the standard curve.

The production of Nitric oxide (NO)/reactive nitrogen species by myoblasts after CTX treatment was measured using the Griess reagent kit (Thermo Fisher Scientific, G7921). Briefly, 100 μL of cell culture supernatant or nitrite standard solution was mixed with 20 μL of Griess reagent and 30 μL of deionized water. The mixture was incubated at room temperature for 30 min and the absorbance of the samples was measured at a wavelength of 548 nm using a plate reader (BioTek Instruments, Synergy H1). The NO concentration was calculated by converting the absorbance readings to the concentration curve of the standard solutions.

### Statistical Analysis

All statistical analyses were conducted using GraphPad Prism 10. The data were analyzed using one-way ANOVA with Tukey’s multiple comparison test and an unpaired t-test. Unless otherwise stated, the results are presented as the mean ± SD. The alpha level was set to 0.05, and differences between the two experimental groups were considered statistically significant at \**p* < 0.05, \*\**p* < 0.01, and \*\*\**p* < 0.001. The symbol "*ns*" represents "no significant difference" between the two groups.

## Acknowledgments

M.Z.-W. gratefully acknowledges the ALIVE initiative (Advanced Engineering with Living Materials) in the SFA-AM program (Strategic Focus Area – Advanced Manufacturing) and the Swiss National Science Foundation Bridge Discovery (40B2-0_211764). P.C. acknowledges funding from the Swiss National Science Foundation Ambizione grant (PZ00P2_216356) and the SNF Spark grant (CRSK-2_220980). O.B.-N. acknowledges financial support from Eccellenza Grant (grant no. PCEGP3_187009) - Swiss National Science Foundation. The Scientific Center for Optical and Electron Microscopy (ScopeM) are thanked for their support on 3D confocal imaging. The authors also acknowledge the Laboratory of Food and Soft Materials at ETH for help with CD measurements. H. L. thanks Jakub Janiak for his kind assistance with the analysis software for muscle contraction.

## Author contributions

Conceptualization: H.L., M.W., A.K.K., I.K., P.C., O.B.-N., M.Z.-W.

Investigation: H.L., M.W., I.K.

Methodology: H.L., M.W., J.J., A.K.K., I.K., P.C., O.B.-N., M.Z.-W.

Visualization: H.L., M.W.

Validation: H.L., M.W., I.K., P.C., O.B.-N., M.Z.-W.

Data analysis: H.L., M.W., P.C.

Software: H.L., J.J.

Supervision: H.L., P.C., O.B.-N., M.Z.-W.

Funding acquisition: P.C., O.B.-N., M.Z.-W.

Project administration: H.L., P.C., O.B.-N., M.Z.-W.

Writing—original draft: H.L., M.W., P.C.

Writing—review and editing: H.L., M.W., P.C., O.B.-N., M.Z.-W. with input from all coauthors.

## Data and materials availability

All data needed to evaluate the conclusions in the paper are present in the paper and/or the Supplementary Materials. Additionally, raw data used in the analyses are available in the ETH Zurich Research Collection (DOI: 10.3929/ethz-b-000710588) under the terms of the repository’s data-sharing policies.

