## Supplementary files for "Filamented Light (FLight) Bioprinting of Mini-Muscles with Self-Renewal Potential"

Hao Liu *et al.*

**This PDF file includes:**

Figures. S1 to S8
Tables S1 to S2

**Other Supplementary Materials for this manuscript include the following:**

Movies S1 to S6

**Video S1**: Spontaneous contraction of myotubes in FLight-engineered mini-muscles (pristine ColFib photoresin) after 2 weeks of differentiation.
**Video S2**: Spontaneous contraction of myotubes observed in bright-field (left) and calcium imaging (right; fluorescence intensity shown in color). Generated from time-lapse confocal microscopy scans.
**Video S3**: Myotube contraction in mini-muscles treated with different molecules, shown in bright-field (left) and calcium imaging (right; fluorescence intensity shown in color).
**Video S4**: Spontaneous contraction of myotubes in mini-muscles biofabricated using pristine ColFib photoresin, before and after CTX treatment.
**Video S5**: Spontaneous contraction of myotubes in mini-muscles created using GelNB photoresin, before and after CTX treatment.
**Video S6**: Spontaneous contraction of myotubes in mini-muscles (pristine ColFib photoresin) after 7 days of recovery from severe CTX-induced injury, treated with different molecules and concentrations.

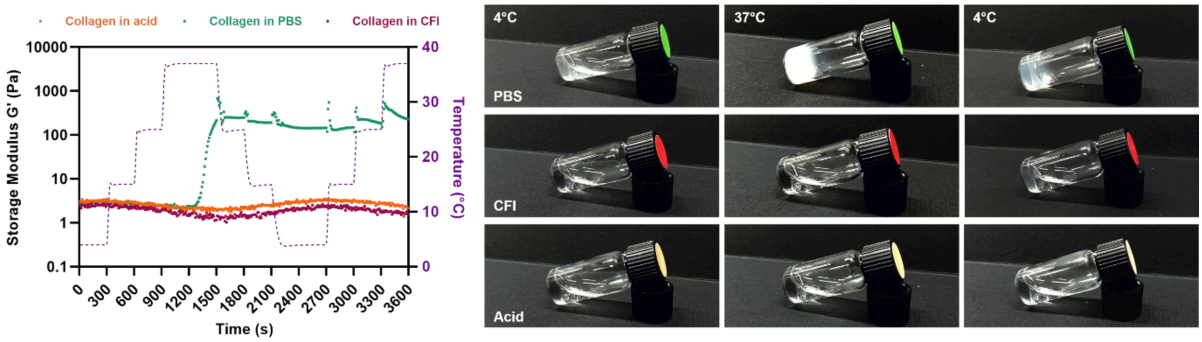


**Figure S1. Thermo-crosslinking of collagen in different solutions.** Rheological analysis of thermo-crosslinking of 5 mg/mL collagen in PBS solution (pH≈7.4), collagen fibrillogenesis inhibition (CFI) solution (pH≈7.2), and in acetic acid (pH≈3.1). Irreversible thermal crosslinking was confirmed in PBS solution but was in CFI or under acidic conditions.

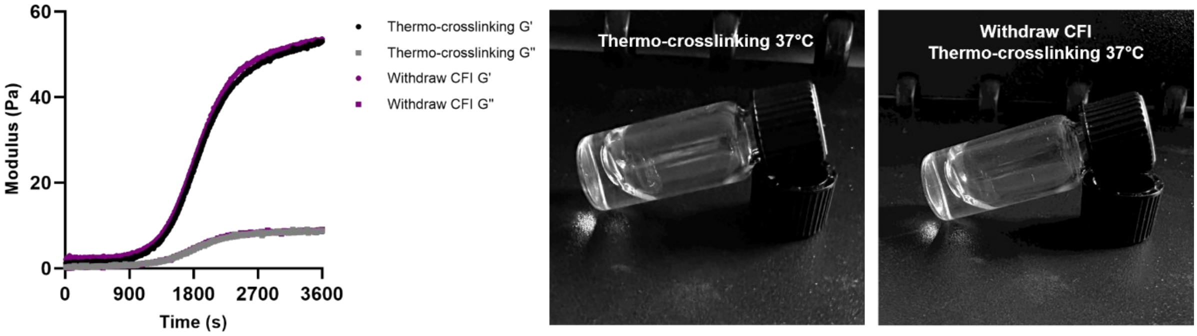


**Figure S2. Thermo-crosslinking of collagen and after removal of CFI solution.** Rheological analysis of thermo-crosslinking of 3 mg/mL collagen in PBS solution (pH≈7.4) and after removal of CFI by dialysis (40 kDa) and redissolved in PBS solution (pH≈7.4). The similar thermo-crosslinking behavior indicates that the inhibition of collagen self-assembly by CFI is transient. The collagen crosslinking at 37°C was still present after removal of CFI solution.

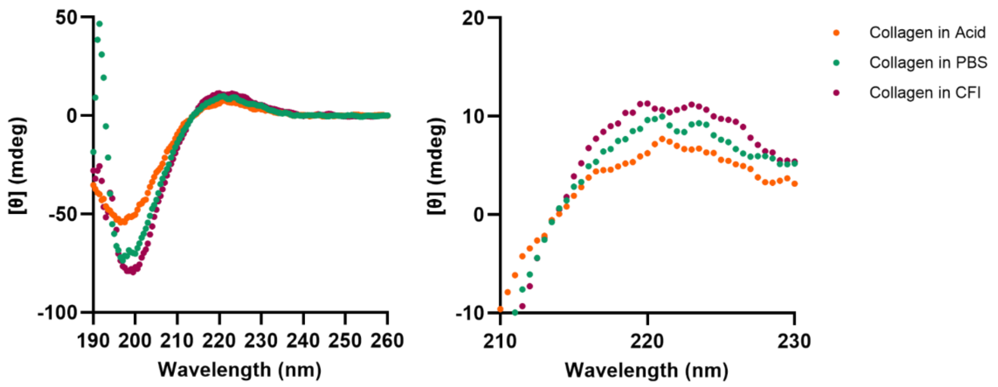


**Figure S3. Circular dichroism (CD) spectra of 0.1 mg/mL collagen type I in different solutions.**

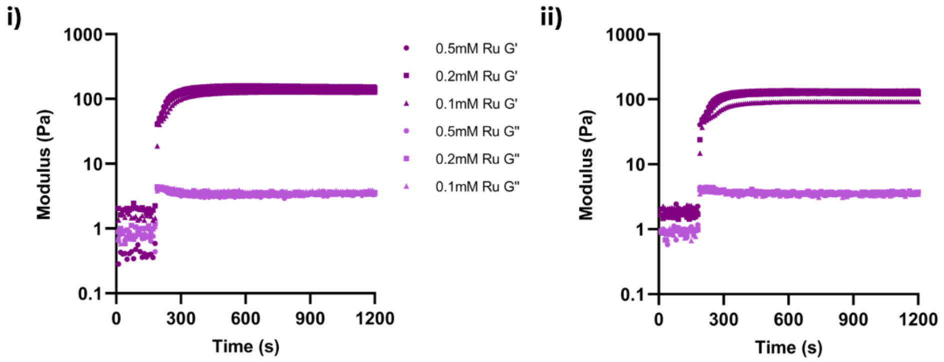


**Figure S4. Photorheological studies of collagen with varying concentrations of photoinitiator.** Rheological analysis of thermo-crosslinking of 3 mg/mL collagen in CFI solution with different concentrations of Ru and the ratio of Ru to SPS. **i**) with different Ru concentrations but a fixed ratio of Ru to SPS (1:10) and **ii**) with different Ru-SPS ratios but a Ru concentration of 0.2 mM.

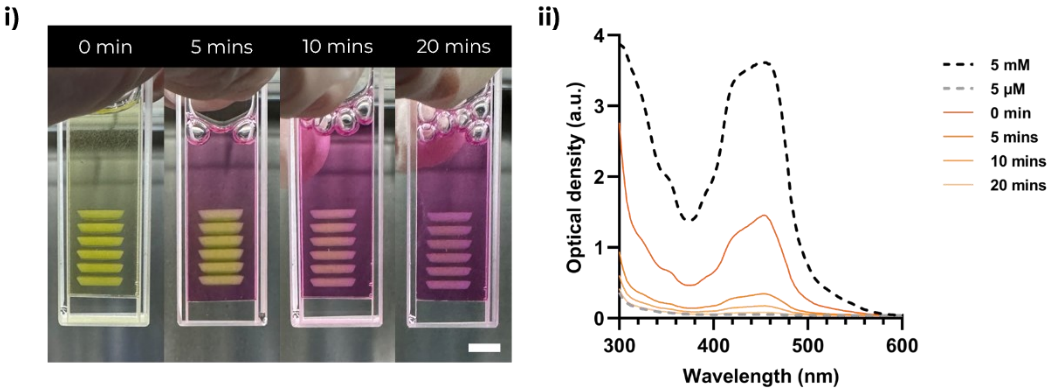


**Figure S5. Removal of ruthenium (Ru) after FLight biofabrication. i**) Photograph of printed FLight muscle constructs with Ru-SPS after 0, 5, 10, and 20 min of washing. Scale bar: 5 mm. **ii**) Optical density of photoabsorption with a wavelength range of 300-600 nm. A reduced absorption peak near 460 nm after 5, 10, and 20 min of washing indicates the decrease of Ru concentration in printed hydrogel constructs. After 20 min of washing, the absorbance approaches that of a standard Ru solution with a concentration of 5 µM.

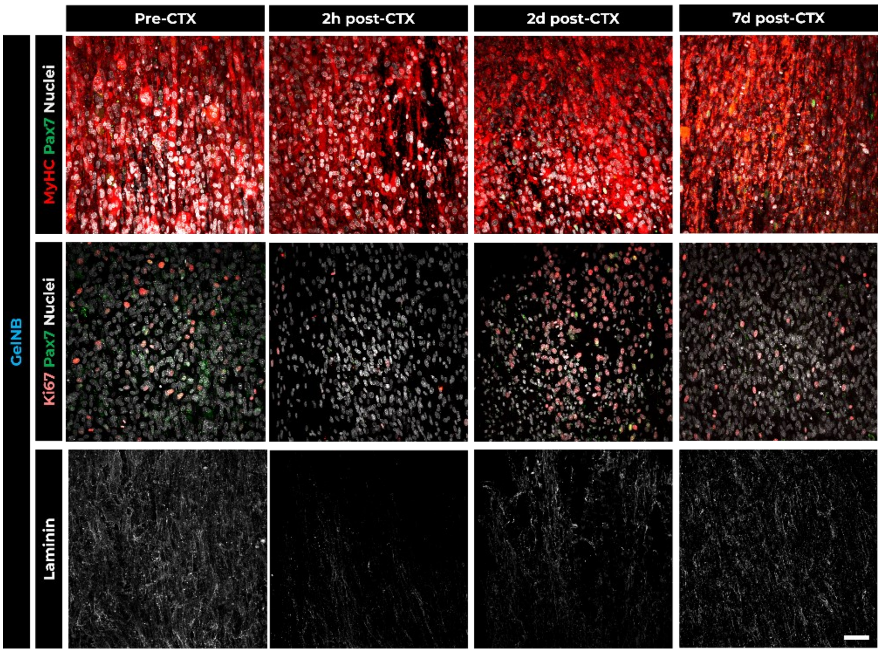


**Figure S6. Self-renewal potential of engineered FLight muscle biofabricated using GelNB photoresin.** Representative confocal images of engineered FLight muscle after 21 days of culture (Pre-CTX), and 2 hours (2h post-CTX), 2 days (2d post-CTX) and 7 days (7d post-CTX) after CTX treatment. Scale bar: 50 µm.

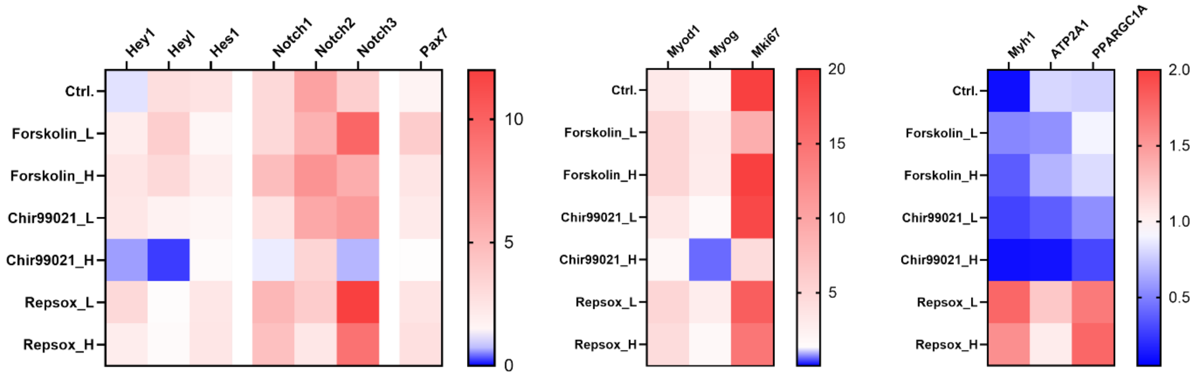


**Figure S7. Gene expression profile of mini-muscles treated with different molecules and concentrations.** Relative change of gene expression in mini-muscles after 7 days of recovery. The fold change shown in the heat map is the mean value. All data were collected from three biological replicates of hydrogel samples with two technical replicates per tissue sample. Details of gene expression are depicted in Figure S8.

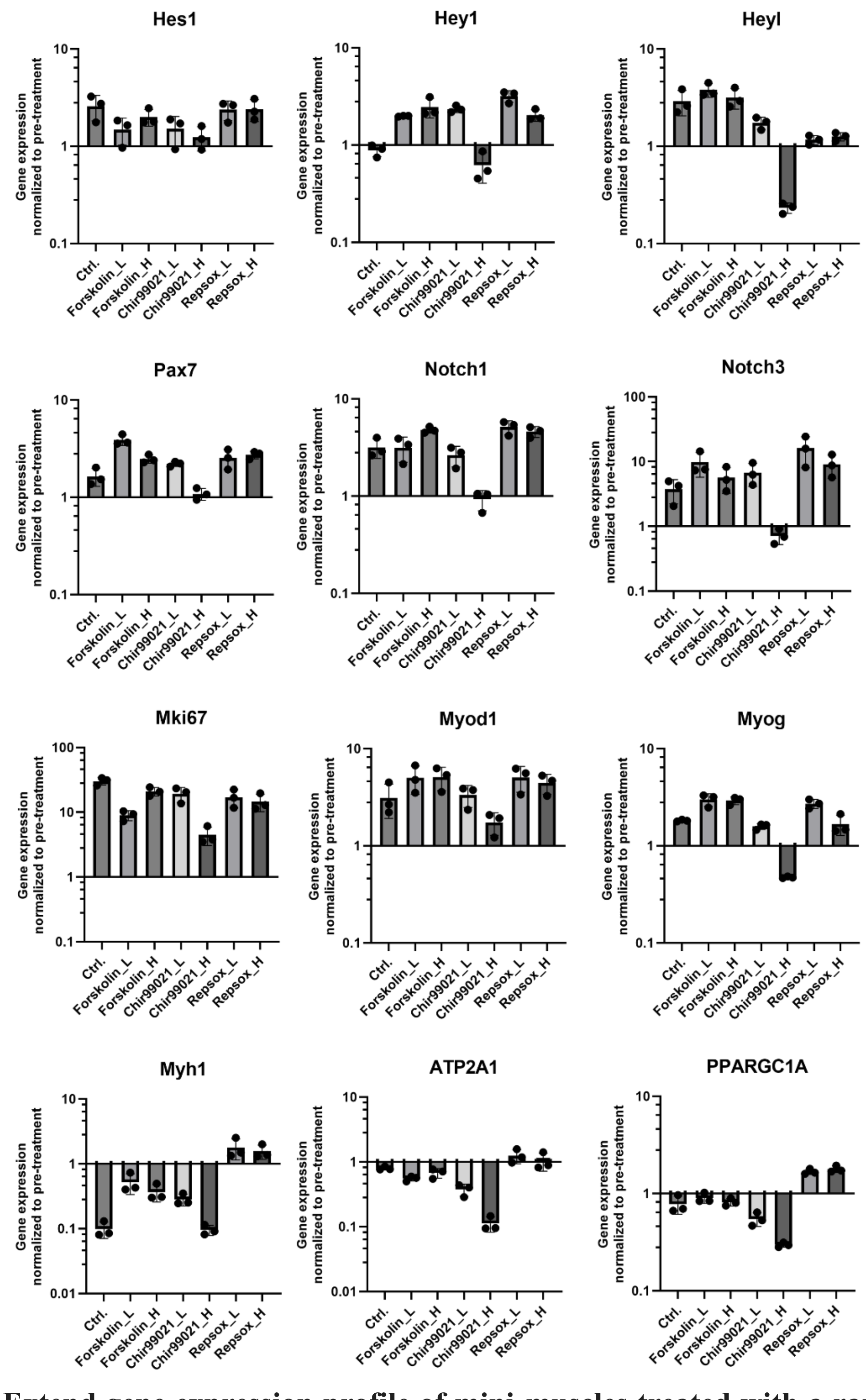


**Figure S8. Extend gene expression profile of mini-muscles treated with a range of small molecules and doses.**

**Table S1: Antibodies and fluorescent dyes used in this study.**

| Resources | Manufacturer | Catalog | Host Species | Dilution |
|---|---|---|---|---|
| **Primary antibodies** | | | | |
| Anti-Ki-67 | BD Biosciences | 550609 | Mouse | 1:200 |
| Anti-MyoD | Invitrogen | PA5-23078 | Rabbit | 1:100 |
| Anti-ITGB1 | Invitrogen | 14-0299-82 | Mouse | 1:100 |
| Anti-MyHC | DSHB | MF20 | Mouse | 1:10 |
| Anti-Sarcomeric alpha actinin | Sigma | A7732 | Mouse | 1:200 |
| Anti-Laminin | Invitrogen | PA1-16730 | Rabbit | 1:200 |
| **Secondary antibodies & Dyes** | | | | |
| Hoechst 33342 | Invitrogen | H3570 | - | 1:1000 |
| Phalloidin-Tetramethylrhodamin B | Sigma | P1951 | - | 1:1000 |
| Goat anti-Mouse IgG(H+L), Alexa 568 | Invitrogen | A-11004 | - | 1:500 |
| Goat anti-Rabbit IgG(H+L), Alexa 647 | Invitrogen | A-21244 | - | 1:500 |

**All antibodies were diluted in 1% BSA-PBS solution if not indicated.*

**Table S2: Sequence of the primer pairs used in the qPCR studies.**

| Gene | Reference | Forward Primer | Reverse Primer |
|---|---|---|---|
| GAPDH | NM_008084.4 | AAGGTCGGTGTGAACGGATTT | GAGGTCAATGAAGGGGTCGT |
| Hes1 | NM_008235.3 | GGTGCTGATAACAGCGGAAT | TTAGGGCTACTTAGTGATCGGT |
| Hey1 | NM_010423.2 | GCCTTTGAGAAGCAGGGATCT | CGTGCGCGTCAAAATAACCT |
| Heyl | NM_013905.3 | AAGAAGCGCAGAGGGATCATA | TTTCTCAAAGGCAGTGGGGAC |
| Pax7 | NM_011039.3 | TCTTACTGCCCACCCACCTA | CCAGGTAATCAACAGCAGTTTGG |
| Notch1 | NM_008714.3 | TGGGCTCCTAACACCTGACT | TGGGGATCAGAGGCCACATA |
| Notch3 | NM_008716.3 | CTCCAGATGCCTGTGAGTCC | CACACTGATGGCCCTGGAAT |
| Mki67 | NM_001081117.2 | ACCCTAGAGGATCTGCCTGG | TCGGGCATCTTTGGGGTTTT |
| Myod1 | NM_010866.2 | GCTCTGATGGCATGATGGATT | ACTGTAGTAGGCGGTGTCGTA |
| Myog | NM_031189.2 | GTGAATGCAACTCCCACAGC | CGCGAGCAAATGATCTCCTG |
| Myh1 | NM_030679.2 | TCCTCATAAAGCTTCAAGTTTGGAC | TATTGGTTGCAGCCCAGTGA |
| ATP2A1 | NM_007504.2 | AAGGCGAAGAAACCGTCACT | GCGTTCTCTGCATTCCGTTC |
| PPARGC1A | NM_008904.3 | GCTGTGTGTCAGAGTGGATTG | AGCAGCACACTCTATGTCACT |
